# Inter- and intra-specific genomic divergence in *Drosophila montana* shows evidence for cold adaptation

**DOI:** 10.1101/282582

**Authors:** Darren J. Parker, R. Axel W. Wiberg, Urmi Trivedi, Venera I. Tyukmaeva, Karim Gharbi, Roger K. Butlin, Anneli Hoikkala, Maaria Kankare, Michael G. Ritchie

**Author notes:** Corresponding authors: Maaria Kankare, Michael Ritchie.

## Abstract

The genomes of species that are ecological specialists will likely contain signatures of genomic adaptation to their niche. However, distinguishing genes related to ecological specialism from other sources of selection and more random changes is a challenge. Here we describe the genome of *Drosophila montana*, which is the most extremely cold-adapted Drosophila species. We use branch tests to identify genes showing accelerated divergence in contrasts between cold- and warm adapted species and identify about 250 genes that show differences, possibly driven by a lower synonymous substitution rate in cold-adapted species. We look for evidence of accelerated divergence between *D. montana* and *D. virilis*, a previously sequenced relative, and do not find strong evidence for divergent selection on coding sequence variation. Divergent genes are involved in a variety of functions, including cuticular and olfactory processes. We also re-sequenced three populations of *D. montana* from its ecological and geographic range. Outlier loci were more likely to be found on the X chromosome and there was a greater than expected overlap between population outliers and those genes implicated in cold adaptation between Drosophila species, implying some continuity of selective process at these different evolutionary scales.

## BACKGROUND

Comparative genomic analyses provide new insights into our understanding of evolutionary processes by helping to identify genes contributing to adaptive divergence (Ellegren 2008; Radwan and Babik 2012). If strong divergent selection due to environmental adaptation or social interactions, such as sexual selection, act as “barrier loci” by influencing species isolation, then identifying them can help to understand the process of speciation (Nosil et al. 2009; Smadja and Butlin 2011). However, accurately identifying such genes is a considerable challenge (Noor and Bennett 2009; Cruickshank and Hahn 2014; Ravinet et al. 2017; Wolf and Ellegren 2017).

Comparative genomic analyses are often hampered by a poor understanding of the sources of selection contributing to species divergence (Ravinet et al. 2017; Wolf and Ellegren 2017). Even when some of the sources of selection seem clear they are often complex and multifaceted, greatly complicating our ability to identify the genetic basis of adaptations. One approach to this problem is to apply comparative genomic techniques to species with distinct ecological specialisations. Several studies have been made of such ecological specialists, including: cactophilic *Drosophila* (Matzkin et al. 2006; Smith et al. 2013), Asian longhorn beetles with specialized feeding habits (McKenna et al. 2016), climate-mediated adaptations in honey bees (Chen et al. 2016), and adaptation to high altitude in humans (Foll et al. 2014). These have successfully identified some associations, but such studies are still relatively few, hindering our general understanding of the genomic landscape of adaptation. Here we describe the genome of *Drosophila montana*, a widely-distributed northern member of the virilis group of *Drosophila*, which shows unique adaptations to seasonally varying environmental conditions prevailing at high latitudes and altitudes. *D. montana* is the most cold-tolerant *Drosophila* species known (Kellermann et al. 2012; Andersen et al. 2015). Their cold tolerance or hardiness involves multiple adaptations, including both a high general resistance to cold and a strong inducible cold acclimation response (Vesala and Hoikkala 2011), as well as a robust photoperiodic diapause (Tyukmaeva et al. 2011), which all contribute to its ability to survive through cold and dark winters. The daily and seasonal activity patterns of *D. montana*, and the interactions and neurochemistry of the core circadian clock genes behind these patterns, differs from those of more temperate species such as *D. melanogaster* (Kauranen et al. 2012; Kauranen et al. 2016; Tapanainen et al. 2018). These features have likely played an important role in allowing *D. montana* to colonize and persist in high-latitude environments (Terhzaz et al. 2015; Kauranen et al. 2016; Menegazzi et al. 2017).

*D. montana* belongs to the *virilis* group of *Drosophila*, which comprises 13 species or subspecies divided into two clades, the virilis and montana phylads, the latter being further split into three lineages (Spicer and Bell 2002). These phylads are thought to have diverged in South Asia during the Early Miocene, after which both of them entered the New World by way of Beringia (Throckmorton 1982). The virilis phylad is constrained mostly within the temperate zone, while the montana phylad has expanded into a variety of habitats and spread to higher latitudes (Throckmorton 1982). Divergence of the two phylads has been estimated to have occurred 7 (Ostrega 1985) to 11 (Spicer and Bell 2002) million years ago, while the North American, European and Asian *D. montana* populations have diverged within the last 450,000 to 900,000 years (Mirol et al. 2007). Interestingly, conspecific *D. montana* populations have been shown to diverge in traits that play a role in ecological adaptation (e.g. Lankinen et al. (2013) and Tyukmaeva et al. (2015)), male sexual cues and female preferences (e.g. Klappert et al. (2007)), and also to show sexual and post-mating pre-zygotic reproductive barriers (Jennings et al. 2014). Information on potential candidate genomic regions and genes for traits involved in cold adaptation and sexual selection has been accumulated through QTL analyses (Schäfer et al. 2010; Tyukmaeva et al. 2015), microarray (Vesala et al. 2012; Salminen et al. 2015), transcriptome (Parker et al. 2015; Kankare et al. 2016; Parker et al. 2016), and RNAi (Vigoder et al. 2016) studies.

Here we aim to identify genes showing evidence of divergent selection linked to cold adaptation by contrasting the genomes of species and populations from different climatic conditions. These analyses were conducted at three levels. Firstly, we classified Drosophila species with well annotated genomes into cold-tolerant and non-cold-tolerant species and used branch tests to identify genes evolving differently between these contrasts. Secondly, we compared *D. montana* with its more temperate relative *D. virilis*. Finally, we compared three divergent populations of *D. montana* from different geographic regions. Such a multilevel approach allows us to identify genes that show recurrent divergence associated with climatic differences between species and populations. Such genes are likely to be particularly important for thermal adaptation, giving insight into the genes and functional processes involved in the evolution of cold tolerance in insects more generally. Thus, our results thus give a novel insight into genomic patterns of selection-driven divergence at different evolutionary scales, in addition to providing a well-annotated genome for a uniquely cold adapted insect species.

## RESULTS

### Genome Sequencing and Assembly

The assembled *D. montana* genome (**Table 1**) has a total length of 183.6 Mb, which falls within the range seen for *Drosophila* species (111-187 Mb), and is similar to that of *D. virilis* (172 Mb), a close relative of *D. montana* with a sequenced genome. CEGMA identified 238 complete orthologs (96%) and 244 partial orthologs (98%) of the 248 CEGMA proteins and BUSCO identified 2457 genes as complete (92%) and failed to identify only 46 (1.7%). RepeatMasker identified that 14.4% of the assembly was composed of repeat elements, the major classes of which were: Simple repeats (4.5%), LTR elements (4.3%), Unclassified (2.9%), and LINEs (1.9%) (**Supplementary Figure 1**). The total percentage of repeat elements identified was around half of that found for related *Drosophila* species *(D. virilis* = 25.9%, *D. mojavensis* = 23.8%, and *D. grimshawi* = 26.1%) likely reflecting the problem of assembling repetitive regions with short reads.

**Table 1.**
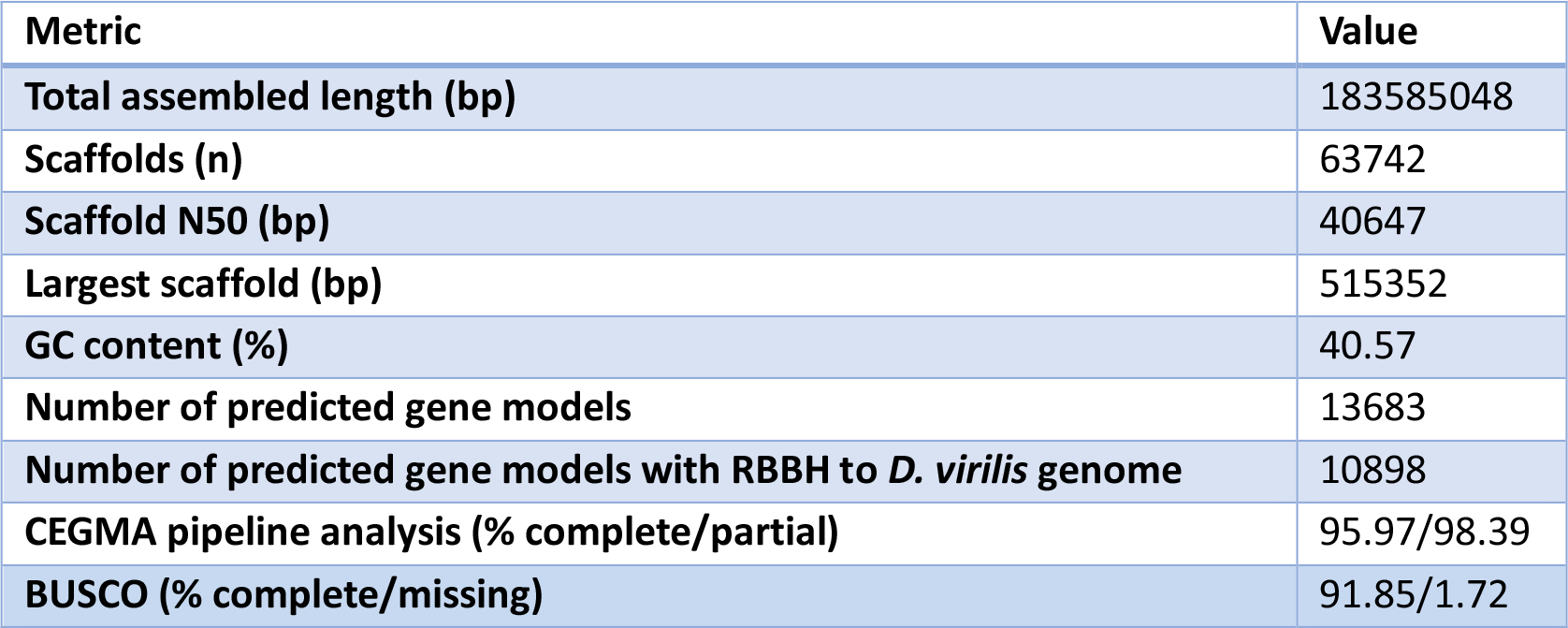
Summary statistics of *D. montana* genome assembly

For the genetic map construction, the final dataset contained 5,858 polymorphic SNPs. The median depth of the SNPs in the final dataset was 52.4 and the average missing data rate was 0.003. The initial analysis formed five major linkage groups (as expected since *D. montana* has five chromosomes in total). Chromosome number was assigned by blasting genes assigned to the linkage groups to the *D. virilis* genome, which have been localised to chromosomes and is largely syntenic with *D. montana* (Schäfer et al. 2010). While the analysis showed clear linkage groups, the order of markers was not totally resolved, likely due to lack of recombination events among F2 progeny (**Supplementary Figure 2**). The tentative scaffold order and position are given in **Supplementary Table 1**. Using this map we were able to anchor approximately one third of the genome assembly to chromosomes. To validate our linkage map we examined coverage of anchored scaffolds. X-linked regions were found to have approximately half the coverage of autosomal regions, as expected since the reference genome was produced from male-only samples (**Supplementary Figure 3**).

### Between-species comparisons identify genes showing accelerated divergence between cold- and warm adapted species

Across the 13 *Drosophila* species we found 250 genes that had significantly different rates of evolution (ω) in cold- and non-cold-tolerant species (**Fig. 1, Supplemental Table 2**). dS was on average lower for cold-tolerant species than for non-cold-tolerant species while dN was very similar (**Fig. 2**, **Supplementary Table 3**). ω was on average greater for cold-tolerant species, probably driven by generally lower values of dS in these species (**Supplementary Table 3**). 203 and 47 genes showed higher values of ω for cold-tolerant and for non-cold-tolerant species, respectively (**Fig. 3**). Genes with elevated ω in cold-tolerant species were enriched for 23 GO terms (Biological Processes: Molecular Functions: Cellular Components = 6:10:7) (FDR < 0.1) (**Supplementary Table 4**), which semantically cluster into the following categories: response to drug, male courtship behaviour, olfaction, ion-channel activity, and developmental processes (**Fig. 4**). Of genes with elevated ω in non-cold-tolerant species we identified 50 enriched GO terms (Biological Processes: Molecular Functions: Cellular Components = 34:3:13) (FDR < 0.1) (**Supplementary Table 5**), which semantically cluster into the following categories: proteasome-mediated ubiquitin-dependent protein catabolic process, reproductive processes, response to fungus, animal organ morphogenesis, regulation of biological and cellular processes (**Fig. 4**). Moreover, DAVID identified 11 functional group clusters for genes with significantly higher ω in cold-tolerant species (**Supplementary Table 6**) including: Nucleotide-binding, Olfaction, Transmembrane proteins, Neural development, Leucine-rich repeat containing proteins, GTPase / GTP binding, Cytoskeleton / Microtubule, and Ion Transport. Finally, DAVID identified 3 functional group clusters for genes with significantly higher ω in cold-tolerant species (**Supplementary Table 7**) including: Calcium-binding EGF domain containing proteins, Transmembrane proteins, and Cytoskeleton / Microtubule.

**Figure 1.**
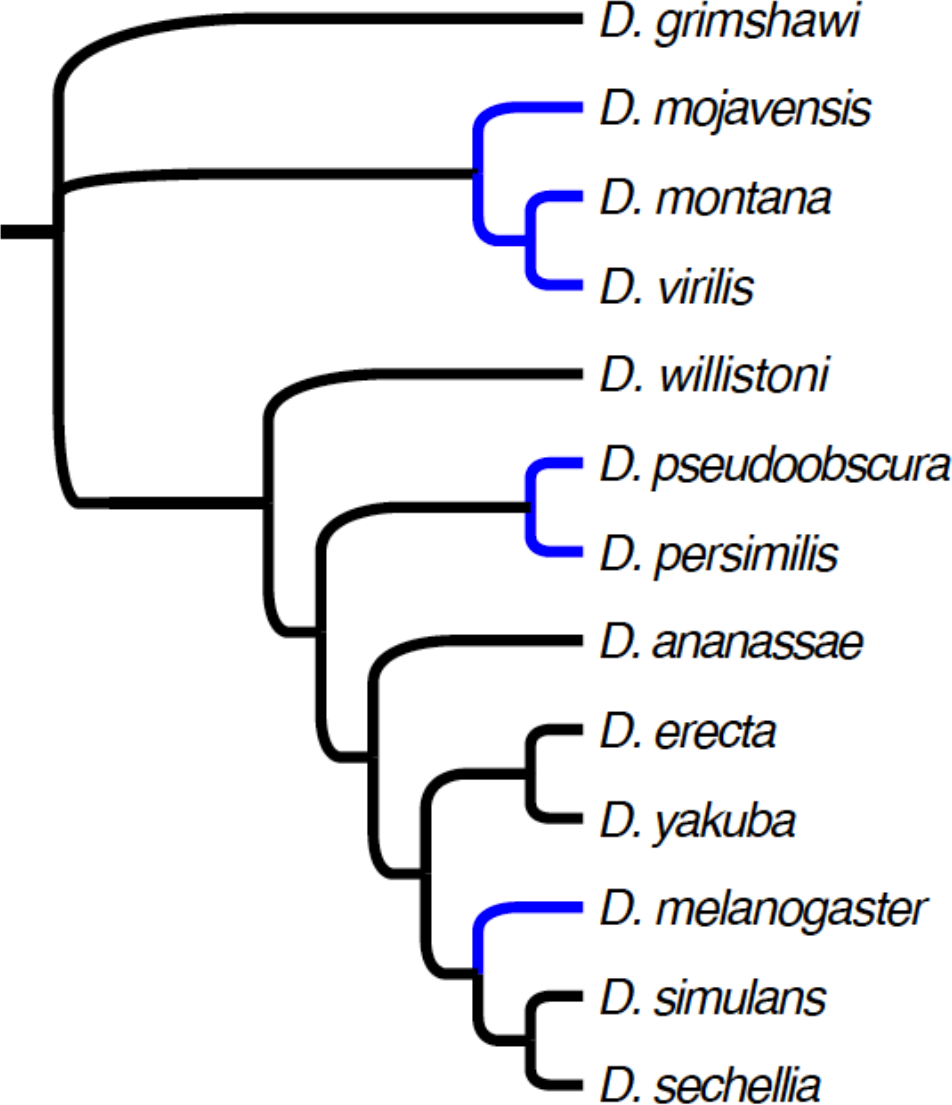
Tree used for multi-species PAML analyses. Cold-tolerant species (species that have a knockdown temperature of <3°C) are shown in blue (data from Kellermann et al. (2012) and MacMillan et al. (2015)).

**Figure 2.**
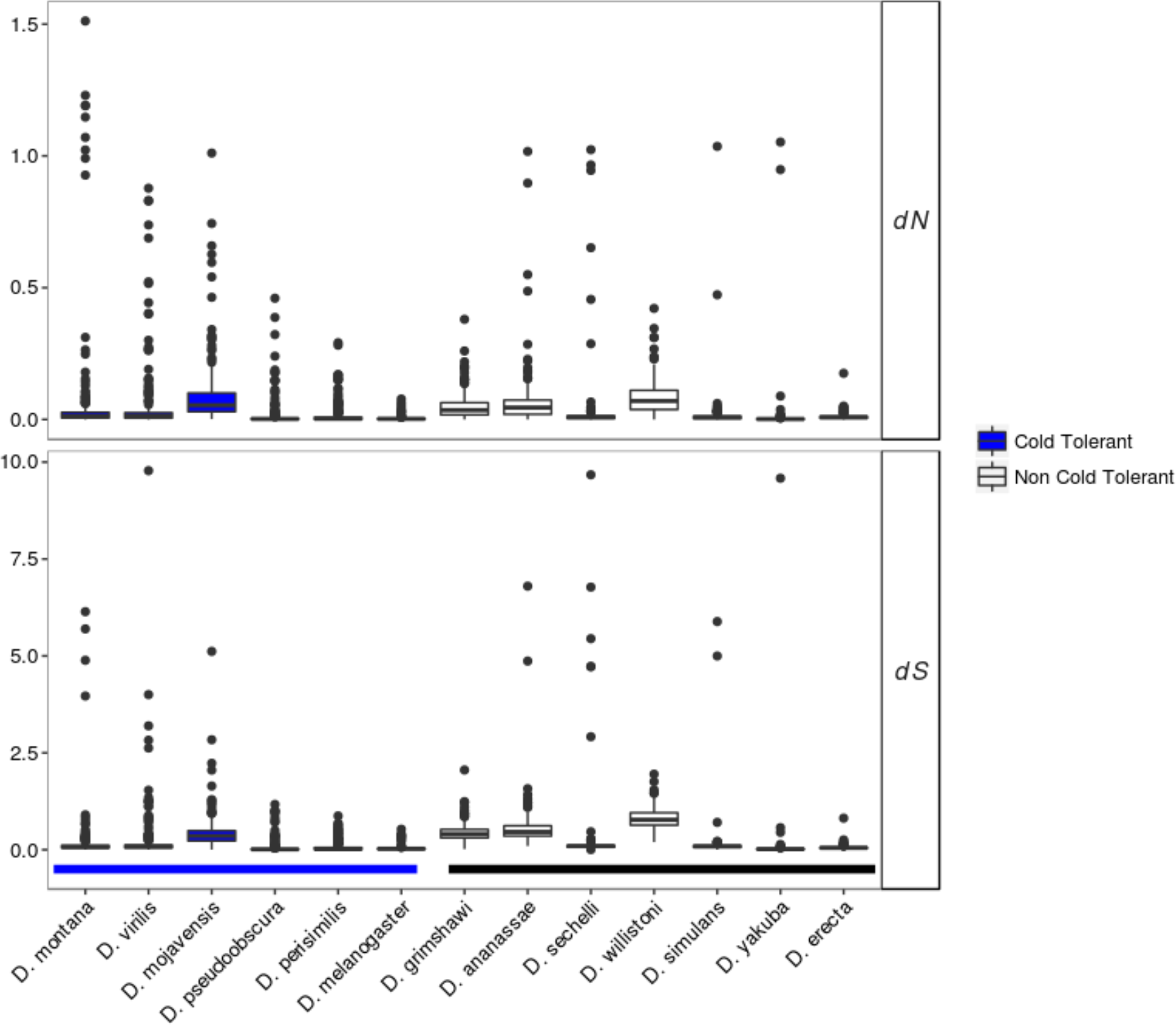
Distributions of dN and dS estimates for each of the 250 genes from 13 *Drosophila* species with significant differences in omega between cold-tolerant and non-cold-tolerant species.

**Figure 3.**
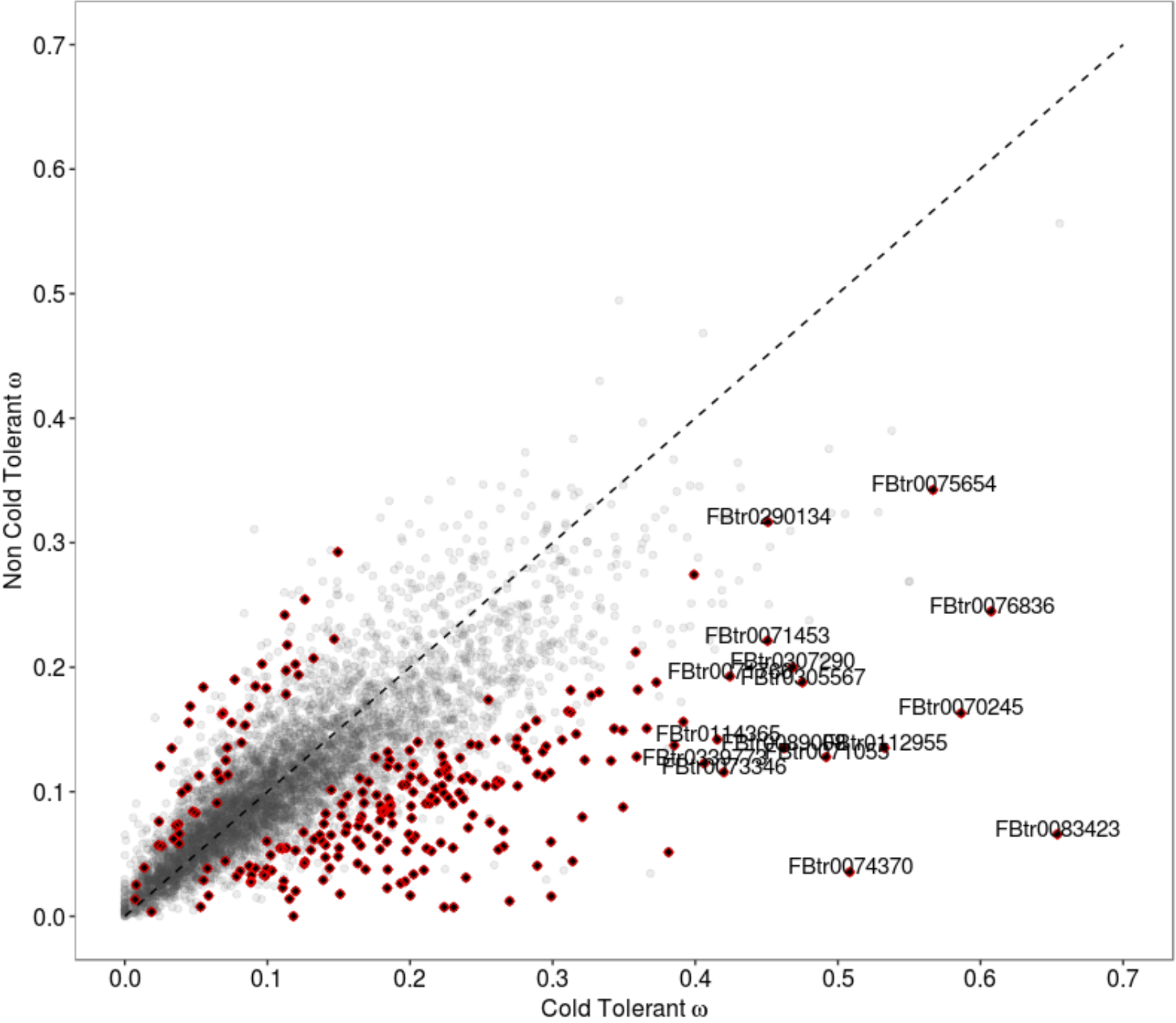
The relationship between values of omega estimated for 5,619 genes in cold-tolerant and non-cold-tolerant species of Drosophila. 250 genes with significantly different estimates of omega are shown in black with red outline. Diagonal line indicates the 1-1 diagonal, points below the diagonal line show elevated levels of omega in cold-tolerant species compared to non-cold-tolerant species, while points above the diagonal show elevated levels of omega in non-cold-tolerant species.

**Figure 4.**
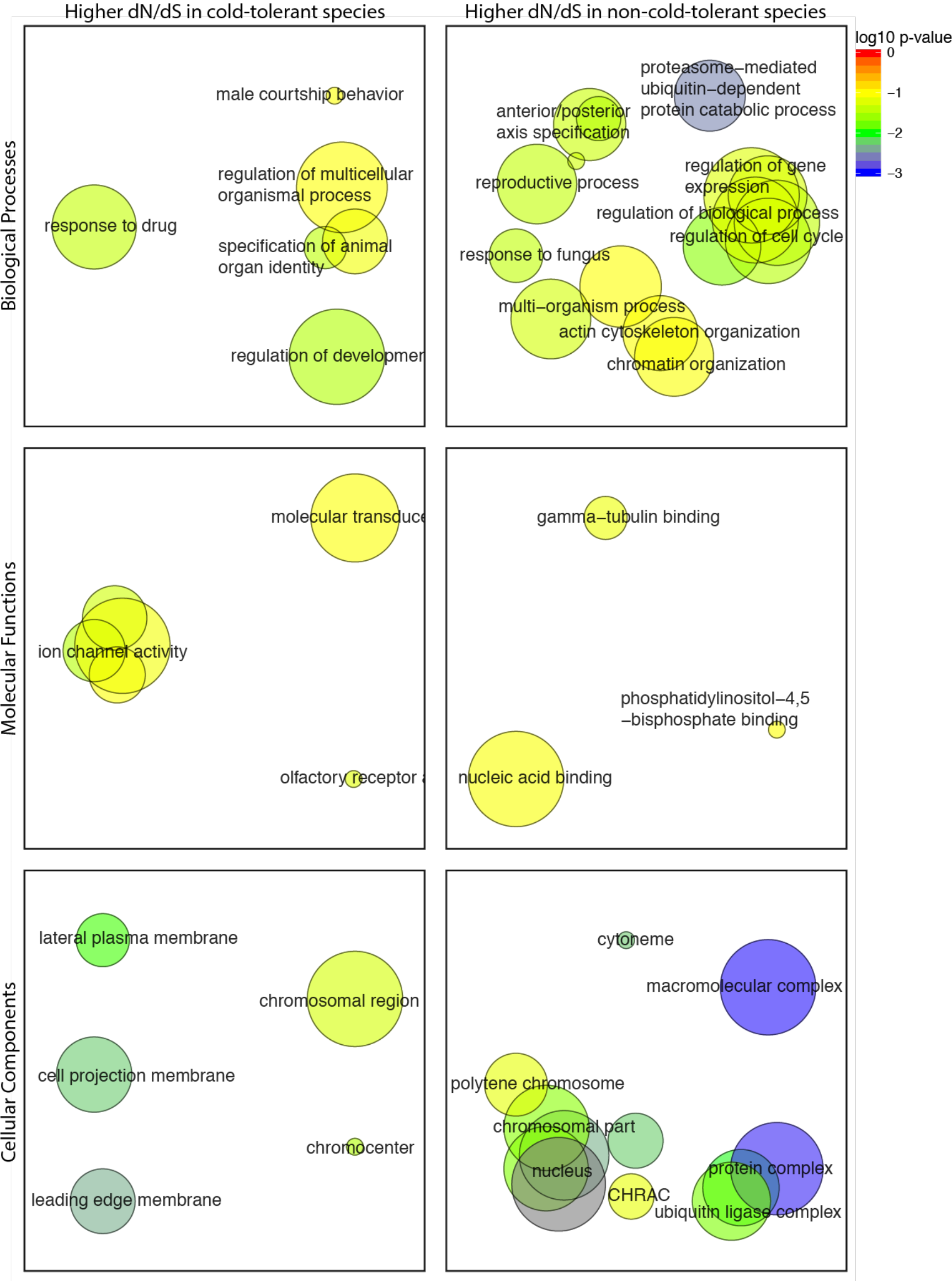
Semantic clustering of significantly (FDR < 0.1) enriched GO-terms for genes showing significantly elevated dN/dS in cold-tolerant or non-cold-tolerant species. Circle size corresponds to the number of genes annotated to the term in the reference database. Circle colour indicates log_10_ FDR of the GO term

### Comparison of *D. montana* and *D. virilis*

We estimated ω (dN/dS) for each of the one-to-one orthologs between *D. montana* and *D. virilis* (**Supplemental Table 2**). No genes had a ω significantly greater than 1 after filtering and multiple-test correction. Comparison of mean ω for several candidate gene sets (genes involved in immune function, reproduction, and cold tolerance) found that none of the candidate genes sets differed significantly from the genomic background (**Fig. 5**). By ranking genes by ω we identified GO terms enriched in genes with relatively high and low ω. For those with high ω we identified 23 enriched GO terms (Biological Processes: Molecular Functions: Cellular Components = 10:4:9) (FDR < 0.1) (**Supplementary Table 8**). Semantic clustering of these GO terms shows that they fall into the following categories: Reproduction, detection of chemical binding / olfaction, amino sugar metabolism, and chitin binding (**Fig. 6**). DAVID identified 9 functional group clusters (**Supplementary Table 9**) including 2 related to chitin production and 2 related to olfactory functions, congruent with the findings from the single GO term enrichment analysis (above). In addition, DAVID also identified 2 clusters involved in: immune defence (C-type lectin domain carrying genes, and Fibrinogen related genes), Transcription factor binding, and a cluster containing genes with either a CAP (cysteine-rich secretory protein) or SCP (Sperm-coating protein) domain. We identified 662 enriched GO terms for genes with low ω between *D. montana* and *D. virilis* (Biological Processes: Molecular Functions: Cellular Components = 485:80:97) (FDR < 0.1). As expected for genes with very low ω the enriched GO terms are consistent with housekeeping roles in the cell (cell cycle control, cell communication, cell developmental process etc.), which are expected to be under strong purifying selection (**Supplementary Table 10**, **Supplementary Figures 4-6**).

**Figure 5.**
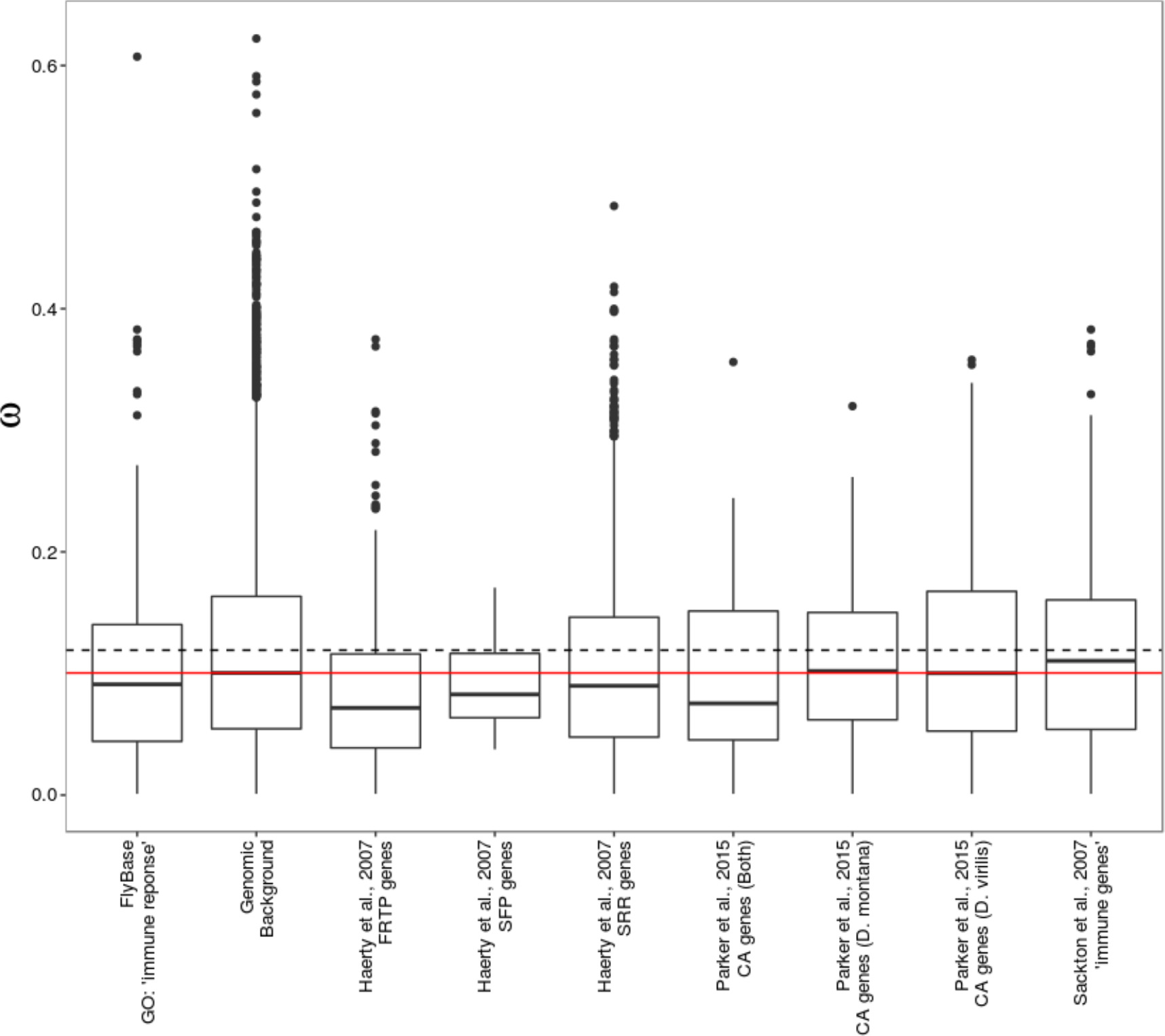
Average values of ω between *D. montana* and *D. virilis* for candidate gene sets. FRTP = female reproductive tract SFP = seminal fluid proteins SRR = sex and reproduction related genes CA = cold acclimation genes. The red and dashed lines indicate the median and mean ω of the genomic background respectively.

**Figure 6.**
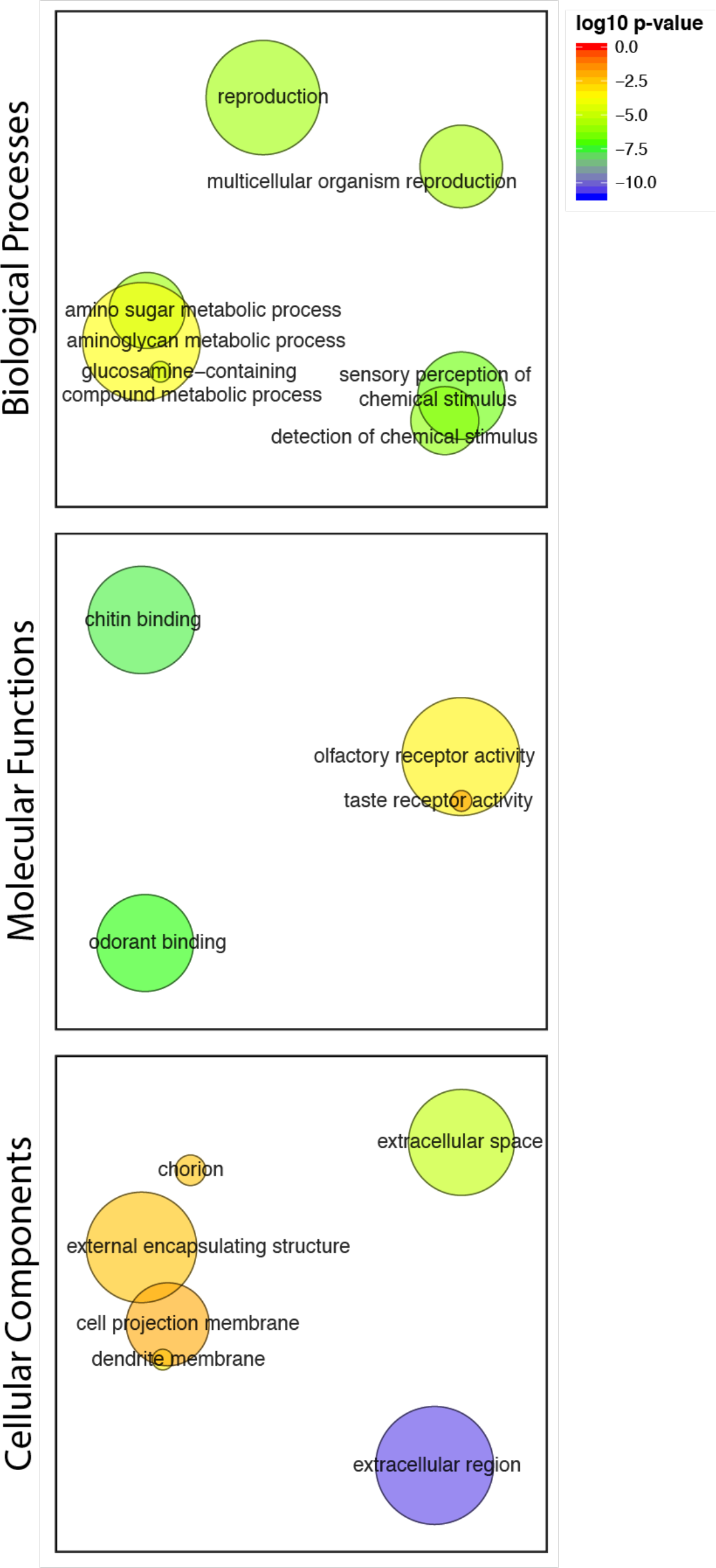
Semantic clustering of significantly (FDR < 0.1) enriched GO-terms for genes showing high dN/dS between *D. montana* and *D. virilis*. Circle size corresponds to the number of genes annotated to the term in the reference database. Circle colour indicates log_10_ FDR of the GO term.

### Genes showing significant between-population divergence are enriched for functional processes associated with cold adaptation

Significant outlier SNPs were found in, or within 1kb of, 1801, 155, and 1387 genes (from pairwise comparisons between Colorado: Oulanka, Colorado: Vancouver, and Oulanka: Vancouver respectively) (see supplementary material for detail on SNP numbers). 10 genes overlapped between all the three pairwise comparisons (**Supplementary Figure 7**, **Supplementary Table 11**). Although this is a relatively small number of genes, it is significantly greater than expected by chance (p = 0.00013). By ranking genes by q-value we could identify GO terms enriched in genes with high divergence for each population comparison (Colorado: Oulanka = 74 (Biological Processes: Molecular Functions: Cellular Components = 27:29:18) (**Supplementary Table 12**), Colorado: Vancouver = 66 (Biological Processes: Molecular Functions: Cellular Components = 19:28:19) (**Supplementary Table 13**), Oulanka: Vancouver = 91 (Biological Processes: Molecular Functions: Cellular Components = 37:39:14) (**Supplementary Table 14**). As with genes, there was a significant overlap of enriched GO-terms between population comparisons (N = 22, p = 1.74 × 10^−79^, **Supplementary Figure 7, Supplementary Table 15**). Semantic clustering of GO terms (**Fig. 7**) and functional clustering (**Supplementary Table 16-18**) showed that the dominant terms include: membrane components, ion transport, small molecule binding, and neuron / synaptic associated terms.

**Figure 7.**
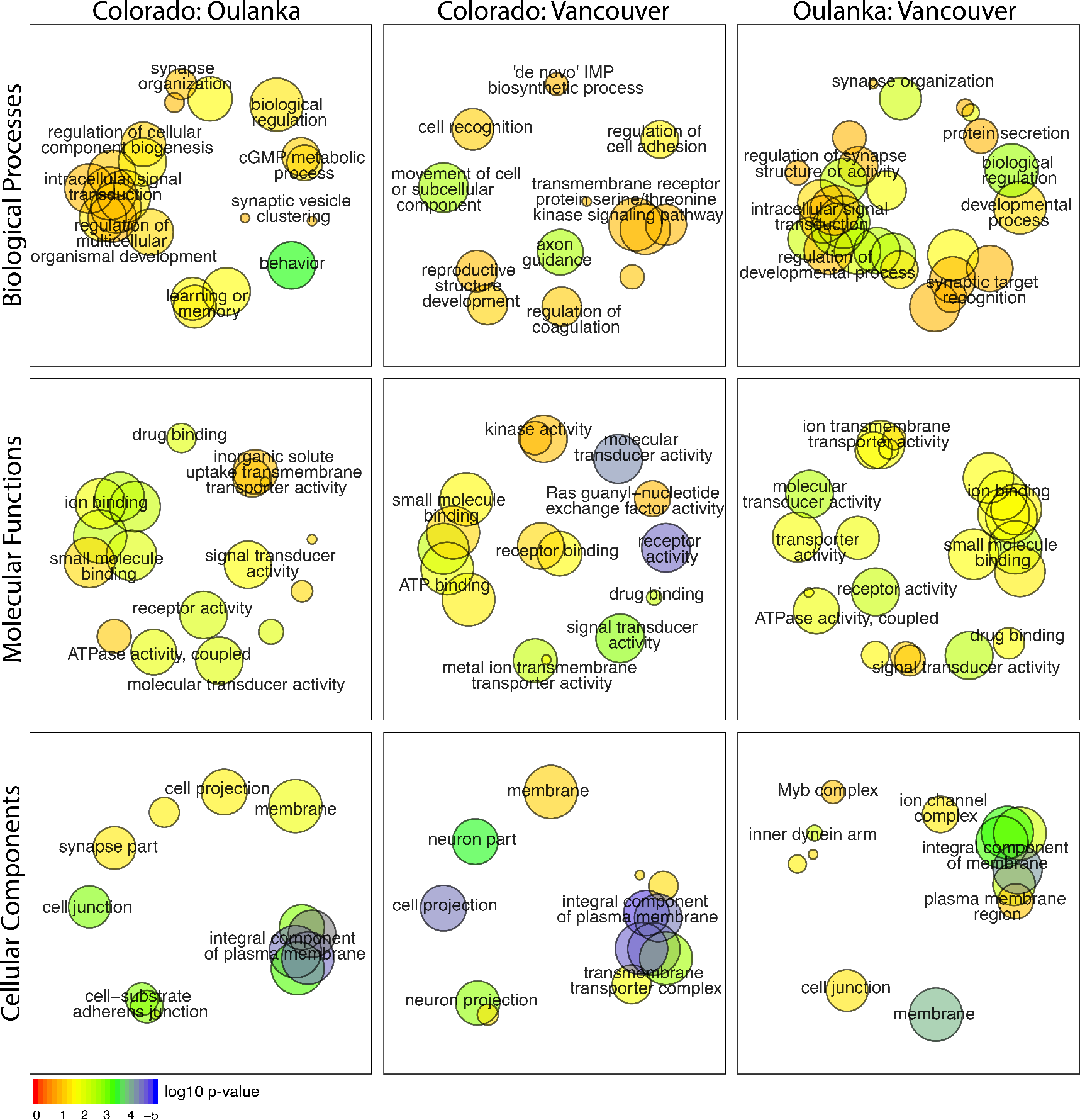
Semantic clustering of significantly (FDR < 0.1) enriched GO-terms for genes showing significant divergence between populations of *D. montana*. Circle size corresponds to the number of genes annotated to the term in the reference database. Circle colour indicates log_10_ FDR of the GO term.

Interestingly, outlier SNPs were not randomly distributed throughout the genome (**Fig. 8 and Supplementary Figure 8**). There was a significant excess of outlier SNPs on the X-chromosome in all pairwise comparisons (Colorado: Oulanka – Chi-squared = 3,029.4, d.f. = 4, p < 0.01; Colorado: Vancouver - Chi-squared = 31.9, d.f. = 4, p < 0.01; Oulanka: Vancouver - Chi-squared = 2477.7, d.f. = 4, p < 0.01). These results held when the proportion of the total genome length of each chromosome was taken to calculate the expected numbers of SNPs.

**Figure 8.**
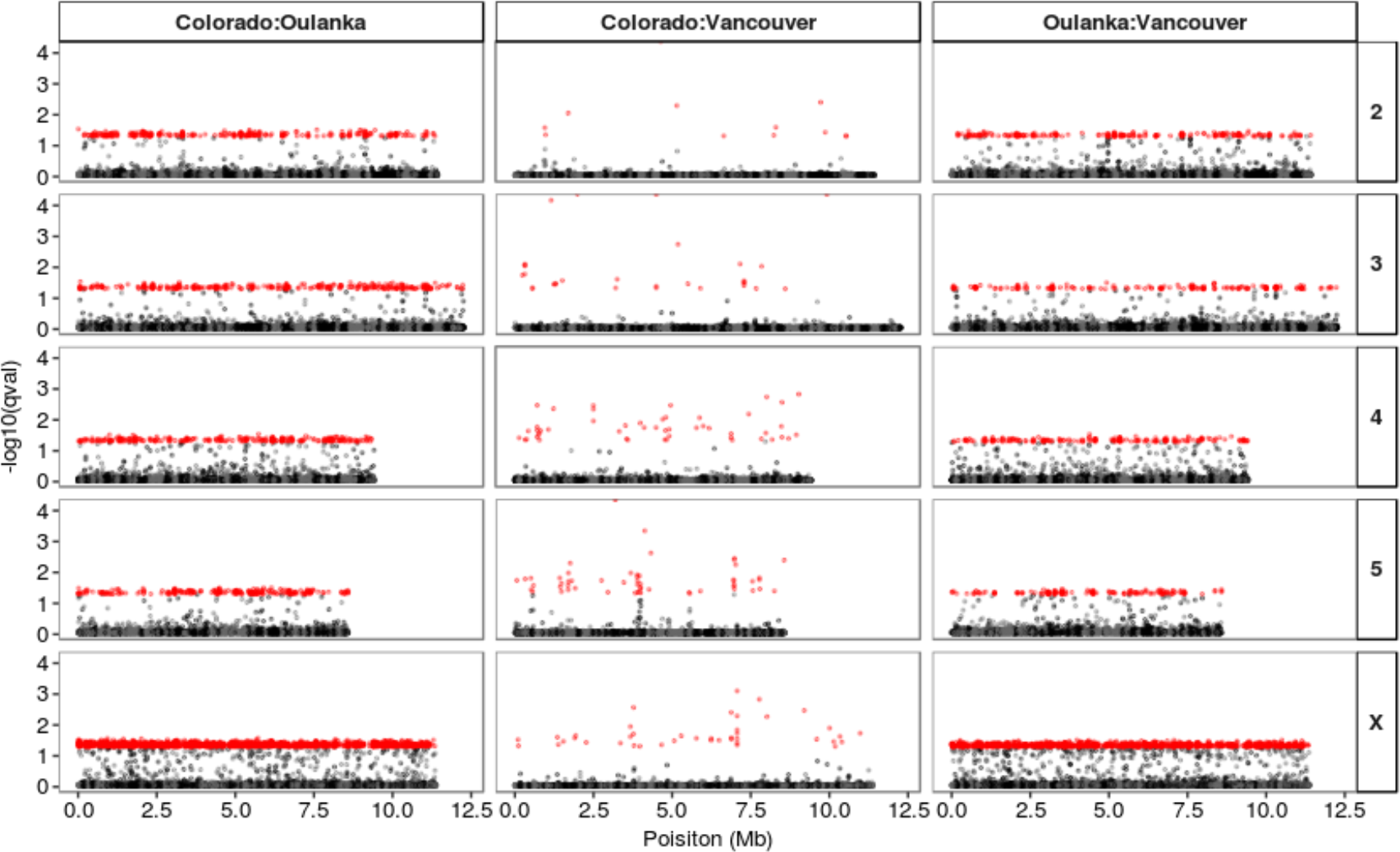
Manhattan plot of q-values from the three pairwise BayeScan analyses for the SNPs on the mapped chromosomes. Red points denote SNPs which passed the 0.05 q-value FDR threshold. Alternating grey and black points denote different scaffolds that have been anchored to the chromosomes. The order of the mapped scaffolds is established but not their orientation.

### Genes showing divergence between species and between populations overlap

We examined whether genes showing significant divergence between populations were the same as those showing higher rates of evolution between cold-tolerant and non-cold-tolerant species. We found 68 genes that had both an elevated rate of evolution between species and significant divergence in at least one population comparison (**Supplementary Table 19**). This is significantly greater than we would expect by chance (Fisher’s exact test = 1.447, p = 0.0006) and implies that genes under divergent selection within species are also more likely to diverge between species. The functions of these genes mirror those enriched in each of the separate comparisons (transmembrane transport / ion transport (9/68), sexual reproduction (16/68), and neurological system process/neurogenesis (15/68)), implicating these genes’ involvement in similar differences in cold adaptation and reproduction between populations and species.

## Discussion

Ecological studies with *Drosophila montana* have shown that it is able to thrive at high latitudes due to a number of adaptations including the evolution of increased cold tolerance and reproductive diapause. By sequencing the genome of this species we were able to use comparative genomics to identify genes and functional processes that differ between *D. montana* and its less cold-adapted relatives. We find evidence for selection acting on neuronal, membrane-transport and ion-transport related genes at both the inter- and intraspecific levels. These findings likely result from selection for an ability to overwinter under harsh environmental conditions, as these processes have clear links to both increased cold tolerance and reproductive diapause.

### Genome assembly and features

We assembled the *D. montana* genome using a combination of Illumina paired-end reads and mate-pair reads. We annotated 13,683 genes, which is comparable to other Drosophila species that have been sequenced (Clark et al. 2007). 10,898 of these genes (80 %) were then assigned to a *D. virilis* ortholog, comparable to the number of orthologs identified between *D. melanogaster* and *D. simulans*. Together with the high BUSCO and CEGMA scores, this suggests that the genic component of the assembled genome is largely complete and successfully annotated.

### Inter- and intra-specific comparisons show evidence for cold adaptation

Firstly, in the comparison between multiple *Drosophila* species, we identified 250 genes with an elevated rate of evolution between cold-tolerant and non-cold-tolerant *Drosophila* species. Interestingly, the increased rate of evolution was biased toward cold-tolerant species, with 77% of these genes showing a higher rate of evolution in these species. Secondly, we compared *D. montana* to its sequenced relative *D. virilis*. Although *D. montana* and *D. virilis* are both relatively cold-tolerant species, *D. montana* is significantly more cold-tolerant than *D. virilis* (Vesala et al. 2012), and *D. montana* is also more desiccation tolerant (Kellerman et al. 2012). In addition, unlike *D. virilis, D. montana* females enter reproductive diapause in late summer, which further increases their chances to survive over the cold season and produce progeny in spring (Watabe 1983). However, genes in the *D. montana* genome showed little evidence for divergent selection when compared to *D. virilis*, most genes showing evidence for purifying selection. Finally, we compared *D. montana* populations from Oulanka, Colorado and Vancouver. These populations face quite different abiotic and biotic conditions throughout the year, and hence can be expected to vary in several traits affecting flies’ life-cycle and stress tolerances. We identified many SNPs that show significant divergence between the three populations; the number of divergent SNPs was smallest between Colorado and Vancouver populations reflecting the likely later divergence times of these populations. Although no divergent SNPs were shared between all population comparisons, when SNPs were grouped by gene, we found evidence for a significant number of overlapping genes. Divergent SNPs were overrepresented in the X chromosome which often shows elevated rates of evolution due to a combination of effects including a smaller effective population size, increased efficacy of selection in hemizygous males, and sexual antagonism. However, as some of the populations are known to differ in sexual behaviour and post-mating pre-zygotic reproductive barriers (Klappert et al. 2007; Jennings et al. 2014), as well as ecological adaptations, it is not possible to distinguish among the multiple possible sources of any divergent selection on X-linked SNPs.

In most of these comparisons the genes with elevated dN/dS or F_ST_ were enriched for functional processes previously demonstrated as important in cold adaptation (see below). In particular changes to membrane components and ion transport, as well as in the neurological system were heavily represented in our enrichment analyses in all comparisons. In addition, we also found enrichment of many small-molecule binding terms, but the specific terms enriched tended to be more varied across the different comparisons. Finally, several comparisons were also enriched for many reproduction-associated terms, which are unlikely to be linked to cold adaptation per se. We discuss each of these functional groups below.

### Functional processes enriched in inter- and intra-specific comparisons

#### Cellular membranes

The composition of the cell membrane is critical for maintenance of cellular function in sub-optimal temperatures (Hazel 1995; Koštál et al. 2003) with changes to cell membrane viscosity shown to be a critical component of cold acclimation in many species (Hazel 1995), including *D. melanogaster* (Cooper et al. 2014). We found enrichment of many terms associated with membrane structure (e.g. intrinsic component of membrane, integral component of membrane, plasma membrane, transmembrane region, etc.) across all our comparisons, providing further evidence for the importance of adjusting cell membrane structure to better survive in cold environments. In addition to these terms, we also found enrichment of other key processes that likely contribute to the functioning of cell membranes at low temperatures. The most important of these are functions associated with cellular ionic balance (e.g. ion channel activity, transmembrane transporter activity, calcium transport, ion binding). Many of the mechanisms involved in the maintenance of cellular ion balance are known to be temperature specific (Heitler et al. 1977; Kivivuori et al. 1990). Failure to maintain the ionic balance of cells leads to metabolic perturbations which can cause a wide range of negative consequences, including cellular damage and even death (Hochachka 1986; Koštál et al. 2004). One class of cells particularly affected by low temperature are neurons (Montgomery and Macdonald 1990; Janssen 1992; Robertson and Money 2012) which are particularly susceptible to cold injury (Hochachka and Somero 2002). In line with this we also found enrichment of several terms related to neuron function (cell projection membrane, dendrite membrane, signal transducer activity, etc.). Finally, we observed that membrane, ion transport, and neuronal terms often functionally clustered together, showing that changes to each of these functions are in fact interrelated. Taken together this suggests that the adjustment of cell membranes for increased cold tolerance is complex, requiring changes to many genes to improve cellular functioning at low temperatures.

#### Small-molecule binding

We observed enrichment of many small-molecule binding terms (small-molecule binding, ATP-binding, kinase, nucleotide-binding, nucleotide phosphate-binding, carbohydrate derivative binding, ribonucleotide binding, anion binding, etc.), both in the population and in the multi-species comparisons. At low temperatures the activity levels of many reactions are reduced meaning that during cold adaptation there is selection to adjust chemical reactions to work better in cold environments (Margesin 2017). In particular ATP-binding and associated terms were enriched in most of our comparisons suggesting that adjustments to ATP-binding may be particularly important for cold adaptation. This finding is supported by the fact that low temperatures adversely affect ATP metabolism across a broad range of taxa (Napolitano and Shain 2004; Morrison and Shain 2008), including freeze-tolerant species like terrestrial earthworm (*Enchytraeus albidus*) that are able to survive winters in a frozen state (Boer et al. 2017).

#### Reproduction

Genes involved in reproduction typically show faster rates of divergence than other genes (Swanson and Vacquier 2002; Clark et al. 2006). Consistent with this we find reproductive-associated terms (male courtship behaviour, single organism reproductive process, reproductive process) are enriched at each comparison level. Different species of Drosophila (including *D. montana* and *D. virilis*) are known to vary for a number of reproductive traits and so this finding is not too surprising. Interestingly, the only pairwise population comparison that shows enrichment for reproductive-associated terms (reproductive structure development, gonad development) is between Colorado and Vancouver. Although all populations show some evidence of reproductive isolation, crosses between Colorado and Vancouver showed the highest proportion of non-developing eggs (Jennings et al. 2014). Moreover, although the exact cause of non-developing eggs is unknown, one possibility is that it could be due to a negative interaction between sperm and the female reproductive tract. Some support for this idea comes from examining the top differentiated genes between Colorado and Vancouver which include the transcription factor *ken and barbie* which has a major role in the development of genitalia of *D. melanogaster* (Lukacsovich et al. 2003).

### Functional processes enriched in specific-comparisons

Although we observe many terms related to cold tolerance common to each of our comparisons (described above), we also observe enrichment of several other functional processes which are restricted to one or two of our comparisons. Of these two (Olfaction and cuticular processes) are of particular interest due to their potential link to cold adaptation and are discussed below.

#### Cuticular related processes

Cuticular and chitin related processes show an extensive enrichment in genes showing elevated dN/dS between *D. montana* and *D. virilis*, but not in the multi-species or population comparisons. Changes to the cuticle are linked to increased cold and desiccation tolerance in insects (Gibbs 2002; Dennis et al. 2015) and in particular to enhancing the stress resistance of the cuticle during diapause (Li and Denlinger 2009; Benoit 2010). This is particularly interesting as *D. montana*, unlike *D. virilis*, has a reproductive diapause meaning the changes we observe in cuticular related genes may have resulted from selection for increased stress resistance to help *D. montana* successfully overwinter. This idea is consistent with the fact that cuticular related processes are only found in the *D. montana – D. virilis* comparison, as this is the only one of our comparisons that directly compares non-diapausing and diapausing capable groups.

#### Olfaction

Drosophila flies have various kinds of olfaction-driven behaviors including the location of food and mates (Amrein 2004; Libert et al. 2007) and the genomic repertoire of olfactory loci is correlated with environmental variation (Gardiner et al. 2008). A cold environment may affect the perception of olfactory signals as the detection of odorants at low temperatures is more difficult due to the reduced concentration of olfactory cues in the air. Previous work in *D. melanogaster* has shown that the sensitivity of the olfactory system increases in response to cold temperature (e.g. Dalton (2000)), and that this change is accompanied by a change in expression in olfactory genes (Riveron et al. 2009; Riveron et al. 2013). Since both sexual and non-sexual olfactory signals are likely to be affected by colder temperatures, we hypothesize that the changes in olfaction-related genes we observe in the present study are a product of adaptation to living in a colder environment as well as of sexual selection to distinguish conspecific flies from the heterospecific ones. Olfaction related terms were enriched in both species-level comparisons, but not population comparisons.

### Population and species divergence at common loci

Phenotypic variation in similar traits between and within species may or may not arise from the same genes even when selection processes are similar (Wittkopp et al. 2009). Here we find that genes which show divergence between populations were also more likely to show elevated differences between species. The functions of these genes mirror those enriched in each of the separate comparisons (transmembrane transport / ion transport sexual reproduction and neurological system processes), implicating these genes’ involvement in similar differences in cold adaptation and reproduction between populations and species. Although any of these genes may be important in cold adaptation, one gene in particular, *Task6*, stands out as an interesting candidate. *Task6* encodes a subunit of two-pore domain potassium (K2P) channels, which are important in setting the membrane potential and input resistance of neurons in Drosophila (Döring et al. 2006). Temperature impacts a cell’s ability to maintain ionic balance, and in particular loss of potassium ion balance has been shown to cause membrane depolarization, induction of chill-coma, and cell death (Andersen et al. 2015; Andersen et al. 2017). As such the changes we observe in *Task6* may be involved in thermal adaptation of species and populations.

## Conclusion

*D. montana* is an exceptional species of Drosophila in terms of cold adaptation, as well as a species used for studies of behavioural variation and reproductive isolation. Here we report the first description of its genome. Although there are few strong signals of divergent selection on coding sequence variation, especially with its closest available relative, contrasts between cold-adapted species and intraspecific population sequencing suggest that the genome contains a clear signal of selection for cold tolerance. We identify many genes potentially important in adaptation and speciation in this ecological specialist species.

## MATERIALS AND METHODS

### Samples and Sequencing

Genomic DNA for the *D. montana* reference genome was extracted from an inbred isofemale line originating from Vancouver, Canada (Can3F9) in summer 2003. This line was inbred via full-sib matings for 37 generations, relaxed for 9 generations and maintained on malt food (Lakovaara 1969) at 19°C in constant light. Quality checked DNA extracted from 210 males using a Gentra Puregene Tissue Kit (Qiagen) was used to produce 3 libraries with different insert sizes: 200bp, 400bp and 3,000bp. The 200bp and 400bp libraries were sequenced using an Illumina HiSeq 2000 at Edinburgh Genomics to produce paired-end reads (101 + 101bp). The 3,000bp library was sequenced using an Illumina MiSeq at The Centre for Genomic Research, University of Liverpool to produce mate-pair reads (101+101bp). This strategy produced 65107854 paired-end reads for the 200bp library, 25618163 paired-end reads for the 400bp library and 19020110 mate-pair reads for the 3000bp library. Reads from the 200bp and 400bp libraries were trimmed using scythe (Buffalo 2014) to remove adaptors and sickle (Joshi and Fass 2011) to quality trim reads (bases with phred quality of <20 were trimmed from the tail end of each read). Reads from the 3,000bp library were trimmed in the same manner, with the addition of a linker sequence removal step.

An initial assembly using reads from the 200bp and 400bp libraries was made using CLC assembly cell (4.0.12). Contigs from this were then blasted (blastN) to two subsets of NCBI’s nt database (arthropod and bacteria) with a e-value threshold of 1 × 10^−40^. Bit scores of blast hits from the arthropod and bacterial databases were compared for each contig, and any with a higher bit score for bacteria than arthropods were considered to be contaminants (**Supplementary Figure 9**). Reads were mapped to contigs identified as contaminants using BWA (v. 0.7.12) (Li and Durbin 2009) and then the unmapped reads were assembled using CLC assembly cell (4.0.12) (default options, minimum contig length = 200bp). Contigs were then scaffolded using the 3,000bp mate pair library using SSPACE-BASIC-2.0. This assembly contained 68950 scaffolds (N50 = 39341). This assembly was then further screened for contaminants using DeconSeq (v. 0.4.3) (Schmieder and Edwards 2011). Bacterial (2786) and viral (4359) genomes were downloaded from NCBI on January 20^th^ 2016 and used as the contamination databases in DeconSeq along with the human genome (hg38). The *D. melanogaster* (r6.09) and *D. virilis* (r1.05) genomes were used as retention databases. DeconSeq identified 5208 scaffolds as contaminants, which were removed from our assembly. We then used this assembly for all subsequent analyses. To assess the completeness of our genome assembly we used the CEGMA analysis pipeline (v. 2.4) (Parra et al. 2007; Parra et al. 2009) which identifies the presence of 248 conserved eukaryote genes, and the BUSCO pipeline (v.1.22) (Simão et al. 2015) which identifies the presence of 2675 conserved arthropod genes.

### Genome Annotation

Full details of the genome annotation are given in the supplementary methods. Briefly, we used the Maker2 pipeline (Holt and Yandell 2011) to first mask putative repeats within the genome, and then used *ab initio* gene predictors SNAP and AUGUSTUS, and gene evidence (from proteins homology and RNA-seq data) to generate gene predictions. Gene predictions from Maker2 were reciprocally blasted to proteins from *D. virilis* (r1.2) with the following cutoffs: e-value < 3 ×10^−13^, query cover > 60% to give reciprocal best blast hits (RBBH). Orthologs for *D. melanogaster, D. sechellia, D. simulans, D. erecta, D. yakuba, D. ananassae, D. persimilis, D. pseudoobscura, D. willistoni, D. mojavensis*, and *D. grimshawi* were then obtained from FlyBase using *D. virilis* FlyBase numbers. Genes without a single ortholog for each species were discarded from multi-species selection analyses (below).

### Linkage map construction

For the genetic map construction, we selected 192 samples from a previous QTL study (Tyukmaeva et al. 2015), which consisted of two families (four parent individuals and their F2 progeny, females only). We used RAPiD Genomics’ (Florida, USA) facilities to develop a set of oligonucleotide probes for 13,975 selected regions in the largest scaffolds of the *D. montana* genome. These probes were used to capture sequence these target loci with 100bp single end reads using an Illumina HiSeq 2000. A resulting SNP dataset was cleaned with Genotype Checker to eliminate possible errors in pedigree/genotyping (Paterson and Law 2011). The R/qtl package (Broman et al. 2003) was used to construct a genetic linkage map after discarding any polymorphic loci that were heterozygous for both parents, duplicated markers, markers showing segregation distortion, and individuals with fewer than 2000 markers. Reads from the 200bp and 400bp genome reference libraries were mapped back to anchored scaffolds using BWA (v. 0.7.12) (Li and Durbin 2009). Multi-mapping reads were discarded. Since the genome reference libraries were produced from males, X linked regions should have half the coverage of autosomal regions, we used the coverage of these scaffolds to validate our linkage map.

## Selection Analyses

### Multispecies analysis

13 species with fully annotated genome sequences available were divided into cold-tolerant and non-cold-tolerant ones; six species with a knockdown temperature <3°C (Kellermann et al. 2012; MacMillan et al. 2015) were classified as cold-tolerant, the remainder as non-cold-tolerant **Fig. 1**). This approach for classifying species was taken to a maximise the power of PAML’s branch tests (see below). To identify genes showing elevated signatures of selection in these species we extracted the longest CDS (N = 5,619) for each ortholog and codon-aligned them using PRANK (v.140110) (Löytynoja and Goldman 2005). Sequences were then analysed in codeml from the PAML (v4.8) package (Yang 1997; Yang and Bielawski 2000). Two models were compared; the “null” model (clock = 0; fix_omega = 0, model = 0, NSSites = 0) which assumes a single common value for ω with an alternative model (clock = 0; fix_omega = 0, model = 2, NSSites = 0) which assumes one value of omega for all the cold-tolerant species and a separate value of omega for the non-cold-tolerant species. Nested models were compared using a likelihood ratio test and p-values corrected for multiple testing using a Bonferroni correction. Additionally, results were filtered to exclude sequences with dN, dS or ω > 10. This comparison tests whether there is a different rate of molecular evolution in cold-tolerant species compared to non-cold-tolerant species.

### Pairwise analysis

To identify protein-coding genes with elevated signatures of selection we estimated pairwise ω (dN/dS) for each gene that had a reciprocal best blast hit (RBBH) to a *D. virilis* gene. The longest coding sequence of each gene and its RBBH ortholog were codon-aligned using PRANK (v.140110) (Löytynoja and Goldman 2005), before estimating ω using codeml in PAML (v. 4.8) (Yang 1997; Yang and Bielawski 2000). To determine if any genes showed ω > 1, we compared genes using a Bayesian estimation of ω in codeml (runmode = −3, model = 0, NSsites = 0) (Angelis et al. 2014) with default priors. The p-values were corrected for multiple testing using a strict Bonferroni correction. We further filtered to exclude any genes where estimates of *dN, dS* or ω were greater than 10.

We then compared mean ω values in several candidate gene sets (genes involved in immune function, reproduction, and cold tolerance) against the genomic background. Genes were classified into two ‘immune’ classes firstly using the GO term ‘immune response’ from FlyBase (version 6.05) and secondly using orthologs of genes identified as being involved in immune function by Sackton et al. (2007). Next, genes connected to reproduction were classified into several reproductive classes following Haerty et al. (2007): sex and reproduction related genes (SRR), female reproductive tract (FRTP) and seminal fluid proteins (SFP). Finally, cold tolerance genes were classified into two classes with genes differentially expressed in response to cold in *D. montana* and in *D. virilis* (Parker et al. 2015). Parker et al. (2015) found that from the differentially expressed genes, 42 were the same in both species but 550 were different, allowing genes to be classified into ‘cold tolerance same’ and ‘cold tolerance different’ groups.

### Population resequencing

For population comparisons we used *D. montana* flies from 3 populations: Oulanka (Finland; 66°N), Crested Butte, Colorado (USA; 39°N) and Vancouver (Canada; 49°N). These populations were established from the progenies of fertilized females collected in the summer of 2008 in Oulanka and Vancouver, and in the summer of 2009 in Colorado. Population cages were set up using 20 F3 generation individuals from approximately 20 isofemale lines for each population. Population cages were maintained at 19°C in constant light (for more details see Jennings et al. (2011)). In March 2013 Genomic DNA was extracted from a pool of 50 females for each population and sequenced at Beijing Genomics Institute using an Illumina HiSeq 2000 to produce paired-end reads (90 + 90bp, insert size = 500bp).

Sequencing produced 84938118 paired-end reads for Colorado and 82663801 for Oulanka. Two runs for Vancouver resulted in 303365095 reads. Reads were quality trimmed (leading or trailing bases with a phred score of <20, or if two consecutive bases had an average phred score of <32 the read was trimmed at this point) and screened for adaptor sequence using trimmomatic (v. 0.30) (Bolger et al. 2014). Reads containing adaptor sequence or that had a length of less than 85 bp after quality trimming, were discarded. Since coverage depth can influence the estimation of allele frequency (Zhu et al. 2012), reads for Vancouver were randomly sampled prior to mapping to the mean number of reads from Colorado and Oulanka. Reads were mapped to the genome assembly using BWA (v. 0.7.12) (Li and Durbin 2009). Reads with a mapping quality of <20 were then removed, and an mpileup file was produced using samtools (v. 0.1.19) (Li et al. 2009). From this, a sync file was produced using PoPoolation2 pipeline (v 1.201) (Kofler et al. 2011). Outlier detection was performed on the raw read count data with BayeScan v. 2.1 (Foll and Gaggiotti 2008; Foll et al. 2010; Fischer et al. 2011), which performs comparably alongside other outlier methods in several simulation studies (Pérez-Figueroa et al. 2010; Vilas et al. 2012; Villemereuil et al. 2014). SNPs were filtered to include only sites with a minimum coverage of 25 and a maximum coverage of 93 (corresponding to the median 10th and 90th percentiles of the population coverage distributions). At the same time, SNPs were only considered if the minor allele had a read count > 4 across all populations. BayeScan was run with 5 pilot runs of 1,000 iterations each followed by a main run of 2,000 iterations, a thinning interval of 10 and a burn in of 1,500. Additionally, three pairwise runs of BayeScan were performed with the same parameters as above. The three pairwise analyses compared Colorado to Vancouver, Vancouver to Oulanka, and Colorado to Oulanka populations, respectively.

### Functional Enrichment

To examine functional enrichment of genes for the species level selection analyses and population level F_ST_ scans, we used GOrilla (Eden et al. 2009). For the pairwise selection analyses genes were ranked by ω (from high to low and low to high). For the multispecies selection analyses, we ranked genes by p-value and direction so that genes with the lowest p-values and a higher ω in cold-tolerant species were at the top, and genes with lowest p-values and higher ω in non-cold-tolerant species were at the bottom, allowing us to identify enriched GO terms for genes showing elevated ω in cold-tolerant species. To examine GO terms for genes showing elevated ω in non-cold-tolerant species the list order was simply reversed. For population level analyses genes were ranked by the most significantly differentiated SNP occurring within 1kb, 10kb, or 100kb of a gene for each population. Results from GOrilla were then visualised using ReviGO (Supek et al. 2011), using the January 2017 version of Gene Ontology.

We used DAVID (v6.8) (Huang et al. 2009a; Huang et al. 2009b) to identify enriched functional groups of genes. A functional group was considered to be significantly enriched if its enrichment score (the geometric mean (in −log scale) of the p-values of the GO terms in the group) was >1 (p < 0.1). For the pairwise selection analyses we identified functional clusters for genes occurring in the top and bottom 10% of genes for ω estimates. For the multispecies selection analyses we identified functional clusters for genes that showed a significantly (FDR < 0.1) higher omega in cold-tolerant species or in non-cold-tolerant species separately. For population level analyses we identified functional clusters for genes containing (within 1kb) significantly differentiated SNPs for each population.

To take advantage of the superior annotation of *D. melanogaster* (Tweedie et al. 2009), we used *D. melanogaster* orthologs for all of the above function enrichment analyses. For the DAVID analyses the ‘background’ list used was the subset of *D. melanogaster* genes available for each analysis.

## Acknowledgements

We would like to thank Paris Veltsos for help with DNA extractions, and RAPiD Genomics (FL, USA) for help with probe design, library preparation, sequencing and initial analysis of the SNP data for the linkage map. This work was supported by the Academy of Finland to AH (projects 132619 and 267244) and to MK (projects 268214 and 272927) and NERC (UK) funding to MGR (grants NE/E015255/1 and NE/J020818/1) and PhD studentship to DJP (NE/I528634/1).

## Data availability

This project has been deposited at NCBI under the BioProject accession PRJNA312336. The accession number for the assembly is LUVX00000000. Raw reads were deposited in the SRA under the following accession numbers: mate-pair reads: SRX1604922, paired-end reads: SRX1602883, SRX1602879, population resequencing reads: SRX1625831, SRX1625832, SRX1625834.

## Authors’ contributions

DJP, AH, MK, and MGR designed the study. KG, UT and RKB conceived the sequencing strategy. DJP and MK performed DNA extractions and original quality checking. DJP, RAWW, UT, and VI, performed the bioinformatics analyses. DJP, AH, MK, and MGR drafted the manuscript, with input from all authors. All authors agreed on the final version of the manuscript.

