## Supplementary Materials for "Inter- and intra-specific genomic divergence in *Drosophila montana* shows evidence for cold adaptation" (from another version of this preprint: version 1, 2018-03-14)

**SUPPLEMENTAL MATERIALS**

**Supplemental methods**

**Genome Annotation**

We used the Maker2 pipeline [1] which first masks putative repeats within a genome and then uses information from *ab initio* gene predictors, and gene evidence (from proteins and RNA) to generate gene predictions. Maker2 uses RepeatMasker (v. 4.0.1) [2] to mask repeats from the Repbase database (downloaded on 8$^{th}$ April 2013) [3]. We also used the default transposable element (TE) sequences from Maker2 to mask TEs.

Maker2 was run in a 2-step iterative process. In the first we provided a trained SNAP model, transcript information, and protein information to Maker2 as described below. To generate a trained SNAP (v. 2006-07-28) gene model, we used the annotation generated from running CEGMA on our genome (above) as a training set. To provide transcript information we used RNA-seq reads from previous studies investigating cold acclimation [4] and diapause [5] (Accession numbers: SRR2910689, SRR2910692, SRR2910693, SRR2910695, SRR2910698, SRR2910701, SRR2910702, SRR2910703, SRR2910704, SRR2910705, SRR1501259, SRR1501260, SRR1501261, SRR1501262, SRR1501263, SRR1501264). These reads were trimmed using SOLiD TRIM (options: -p 4 -q 25 -y on -e 3 -d 10) [6] and the reads that passed this filter were then error corrected using SOLiD Accuracy Enhancer Tools (SAET) to reduce the amount of color calling errors, or erroneous bases, in the sequence. These reads were then mapped to the genome assembly with TopHat2 [7] (v. 2.0.10) (options: --read-mismatches 3 --read-gap-length 3 --read-edit-dist 3) and Cufflinks [8] (v. 2.1.1) (default options) to produce an annotation (gtf) file. This file was converted to GFF3 format using the cufflinks2gff3 script so it could be passed to Maker2. Protein homology evidence was provided using protein sequences from FlyBase for *D. virilis* (r1.2) and *D. melanogaster* (r5.56) and Maker2 was run using default options except for est2genome=1, protein2genome=1. The annotations from this round were then used to train the SNAP and AUGUSTUS (v. 2.5.5) gene predictors and Maker2 was run again using the protein and transcript evidence as before, the updated SNAP and AUGUSTUS gene models and default options except est2genome=0, protein2genome=0. Maker2 calculates the Annotation Edit Distance (AED) for its annotations which quantifies how well predictors match evidence and we used this score to select annotations with AED < 1.
