## Supplementary Materials for "Inter- and intra-specific genomic divergence in *Drosophila montana* shows evidence for cold adaptation" (from another version of this preprint: version 1, 2018-03-14)

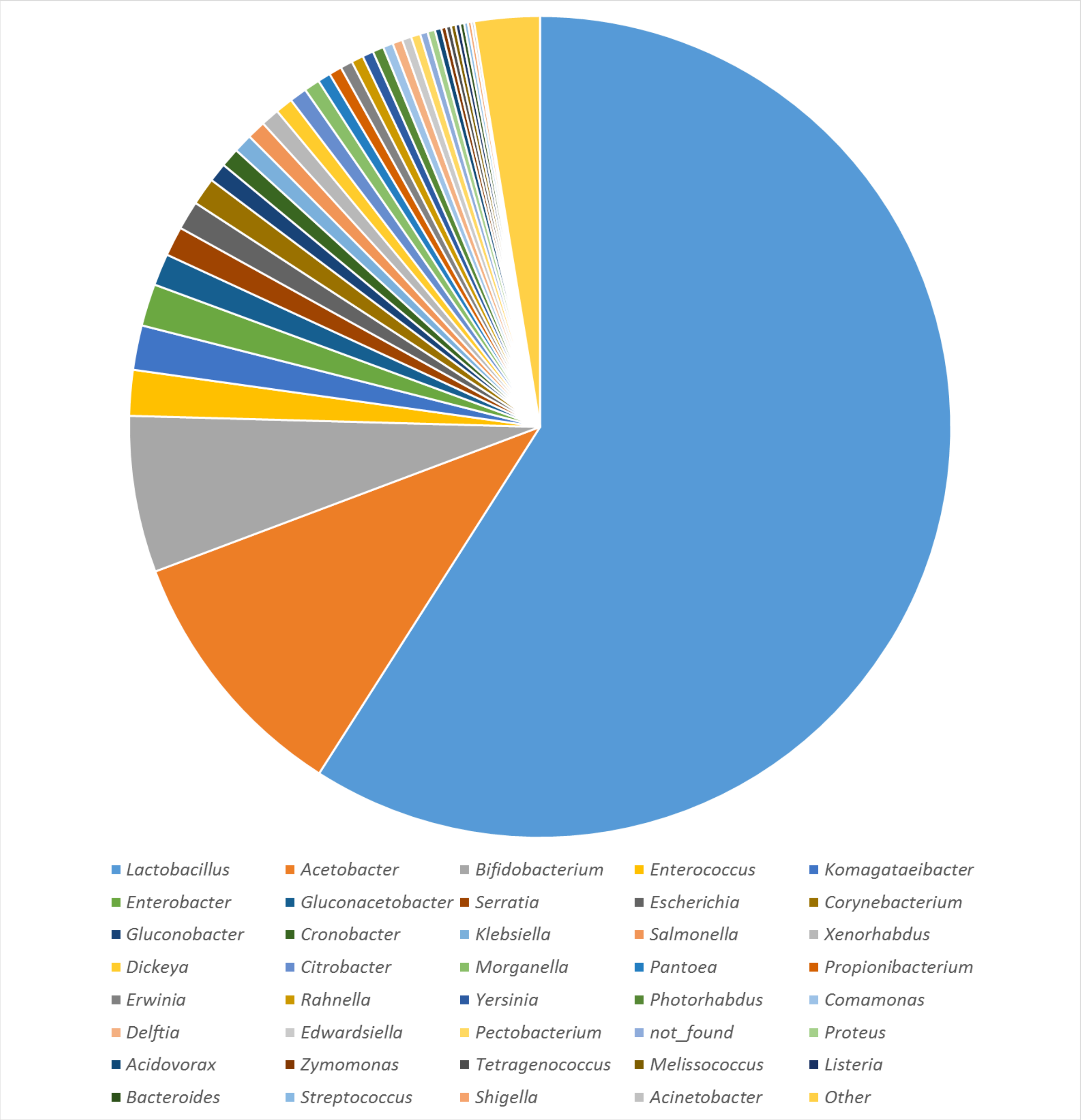


**Supplemental Figure 1 |** Species of blasts hits of scaffolds classed as contaminants

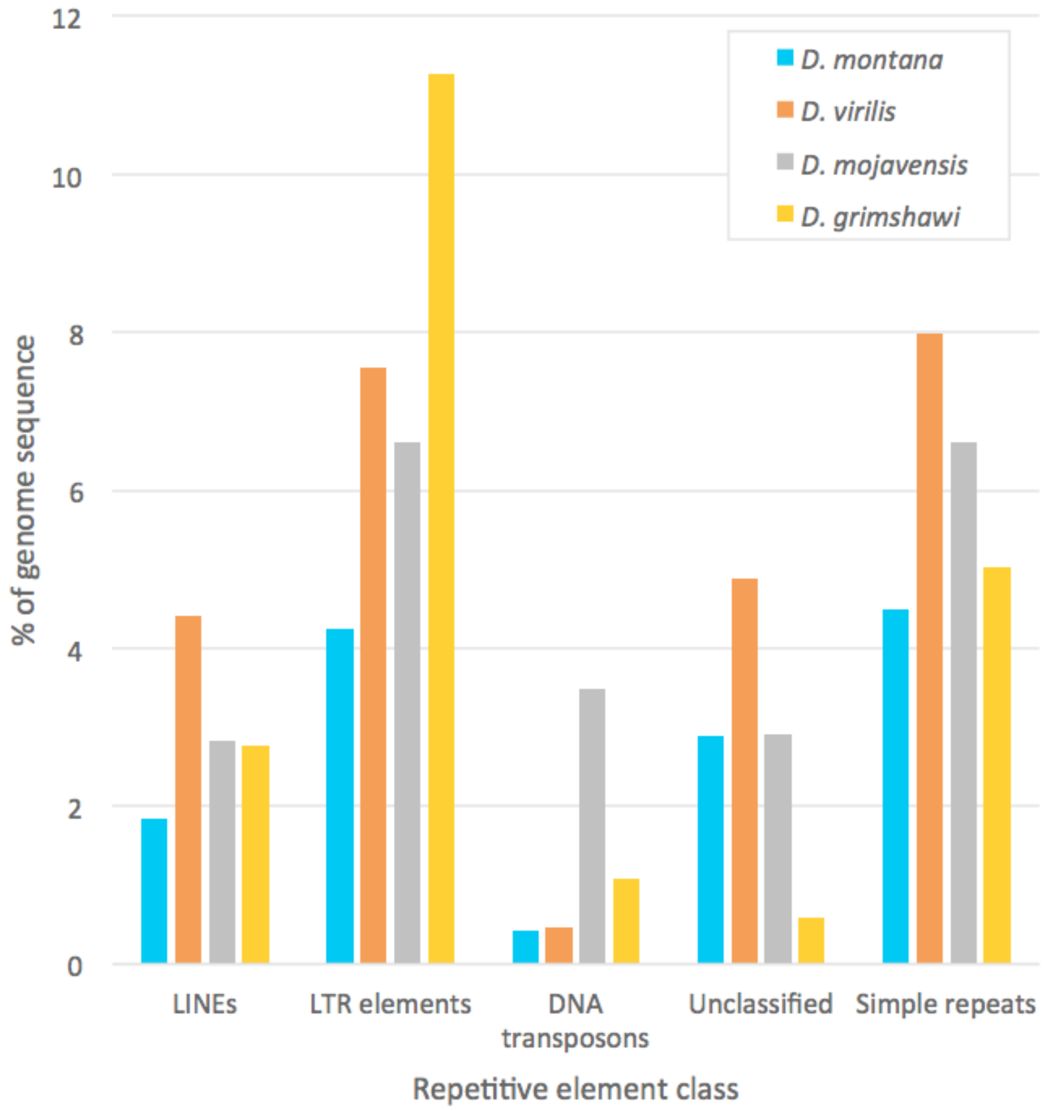


**Supplemental Figure 2 |** Major repetitive elements classes in *D. virilis*, *D. montana*, *D. mojavensis*, and *D. grimshawi*.

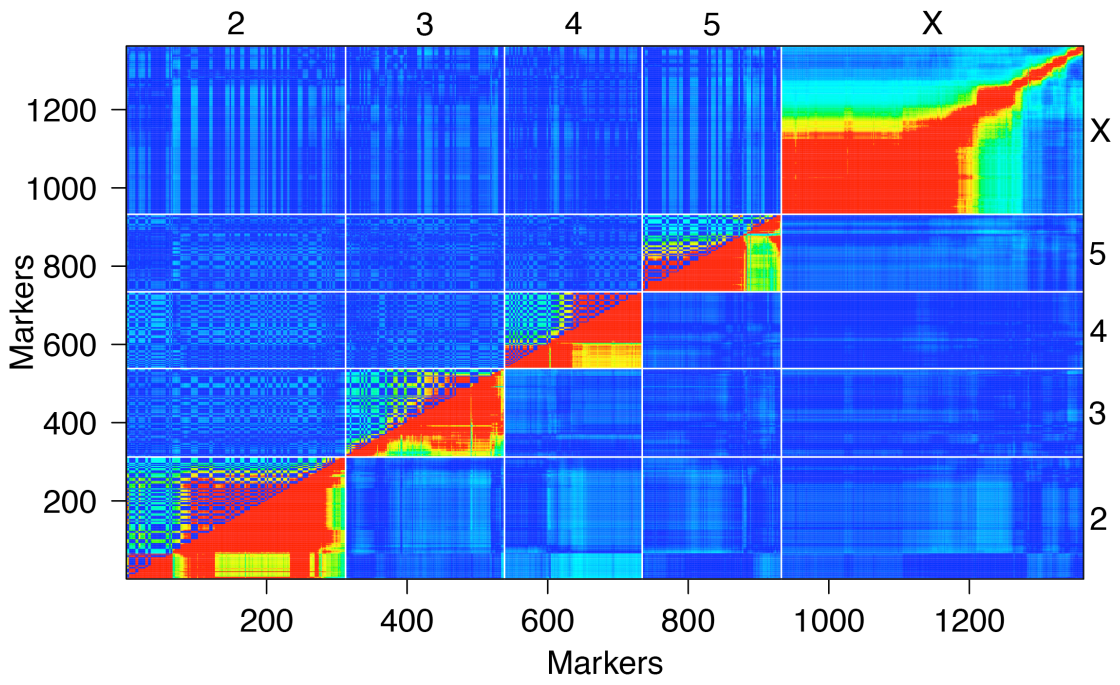


**Supplemental Figure 3 |** Plot of estimated recombination fractions (upper-left triangle) and LOD scores (lower-right triangle) for all pairs of markers. Red indicates linked (large LOD score or small recombination fraction) and blue indicates not linked (small LOD score or large recombination fraction).

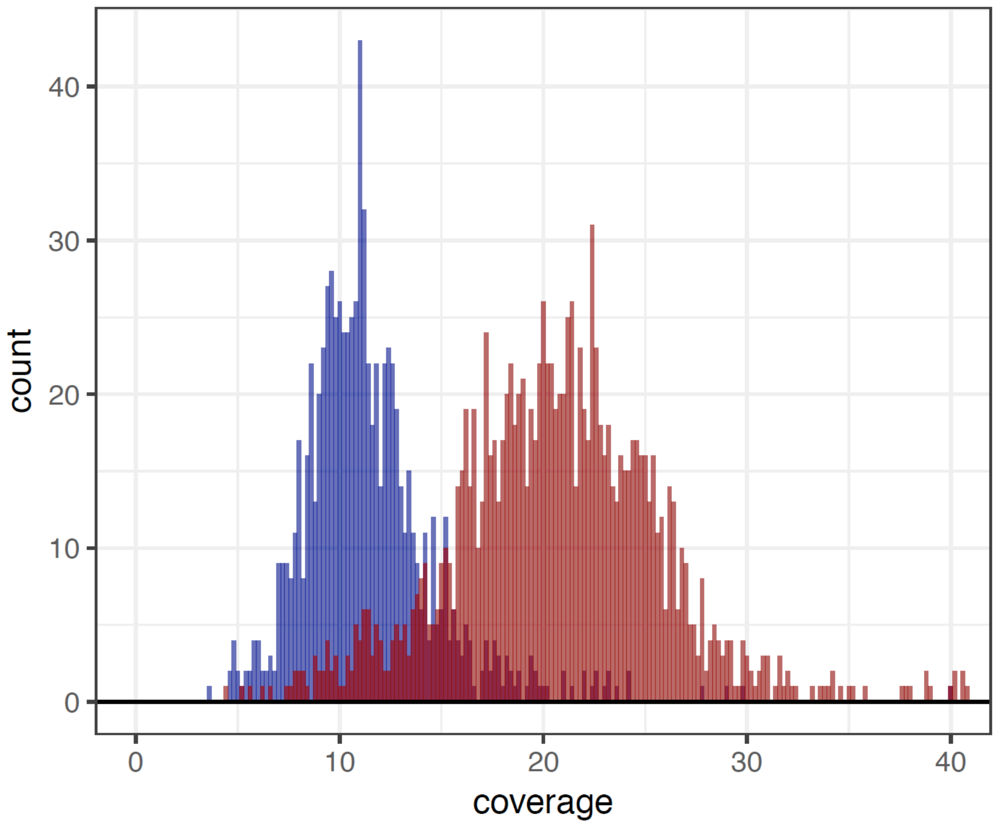


**Supplemental Figure 4 |** Coverage of contigs in the linkage map. Blue are those identified as sex-linked from the linkage map, red as autosomal.

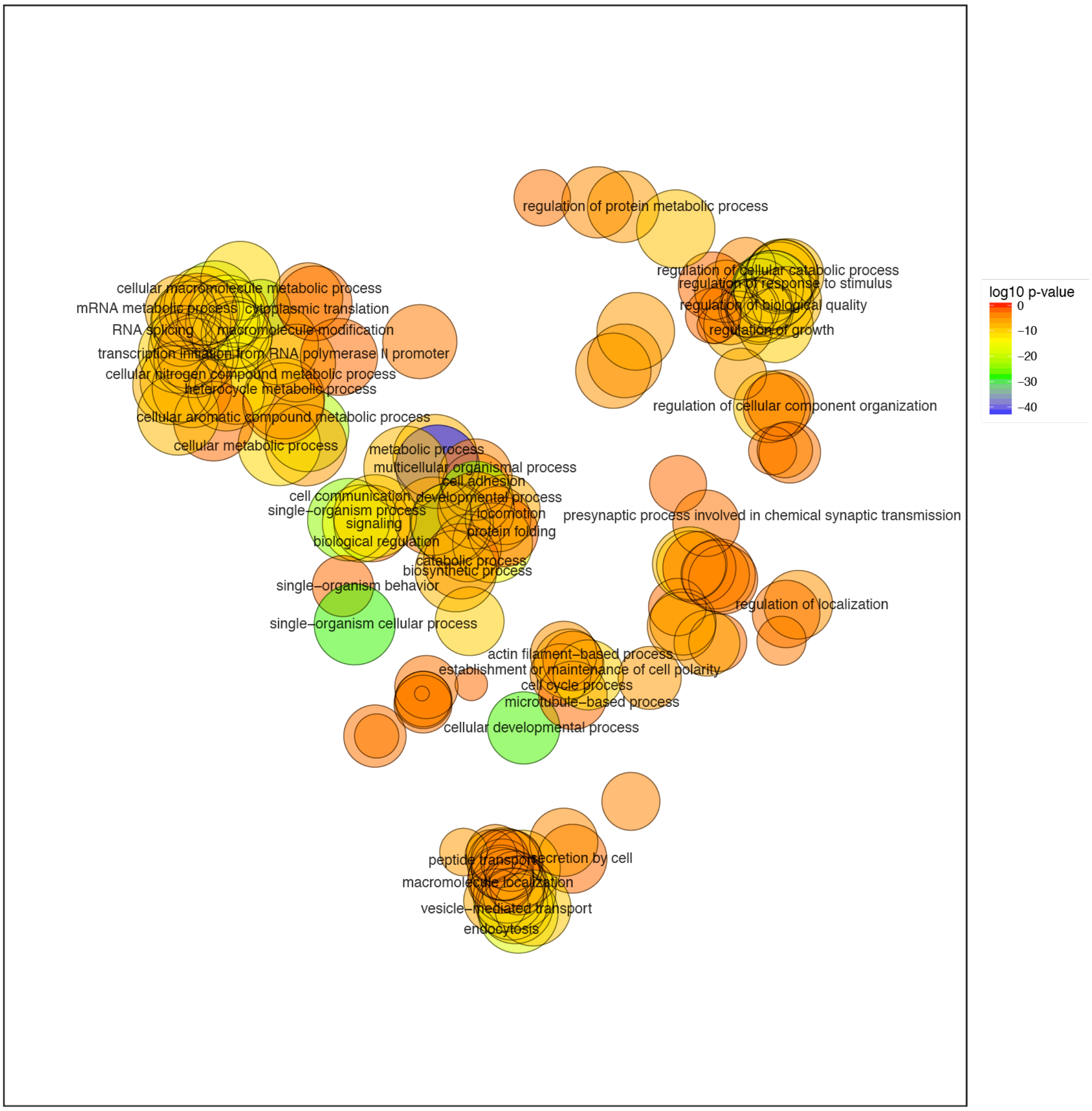


**Supplemental Figure 5** | Significantly (FDR < 0.1) enriched biological process GO-terms for genes showing low dN/dS between *D. montana* and *D. virilis.* Circle size corresponds to the number of genes annotated to the term in the reference database. Circle colour indicates log10 FDR of the GO term

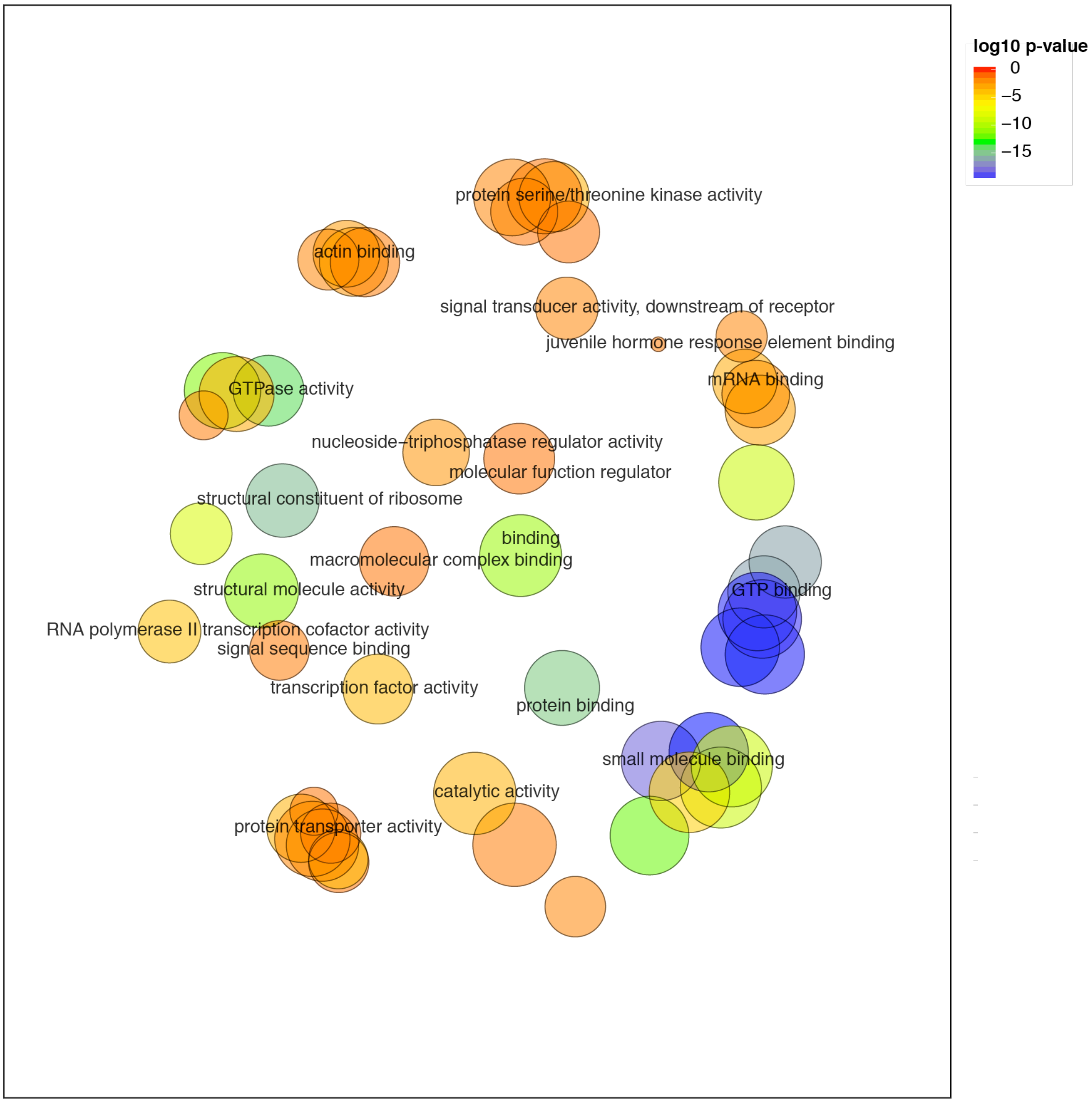


**Supplemental Figure 6 |** Significantly (FDR < 0.1) enriched molecular function GO-terms for genes showing low dN/dS between *D. montana* and *D. virilis.* Circle size corresponds to the number of genes annotated to the term in the reference database. Circle colour indicates log10 FDR of the GO term

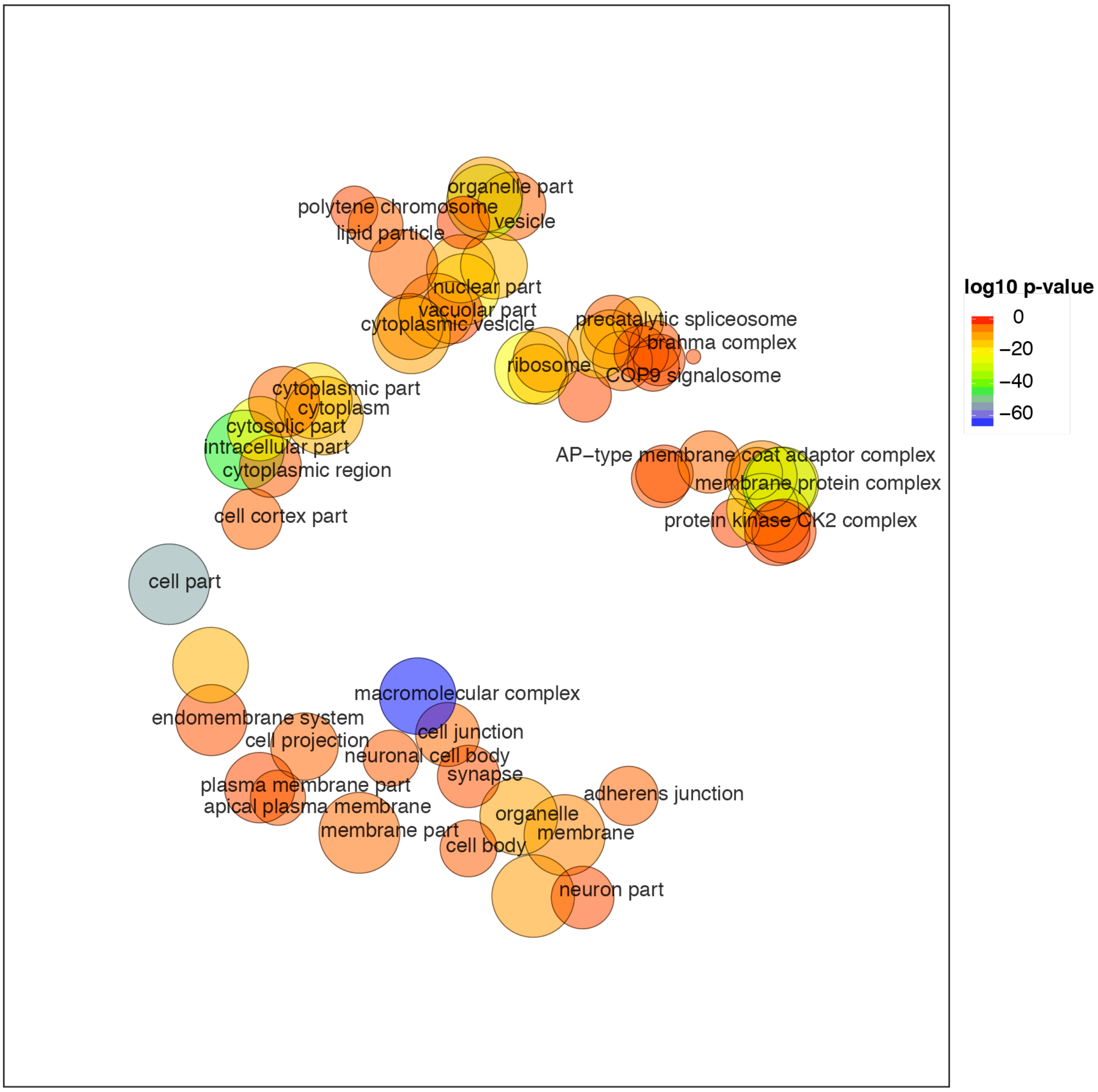


**Supplemental Figure 7 |** Significantly (FDR < 0.1) enriched cellular component GO-terms for genes showing low dN/dS between *D. montana* and *D. virilis.* Circle size corresponds to the number of genes annotated to the term in the reference database. Circle colour indicates log10 FDR of the GO term

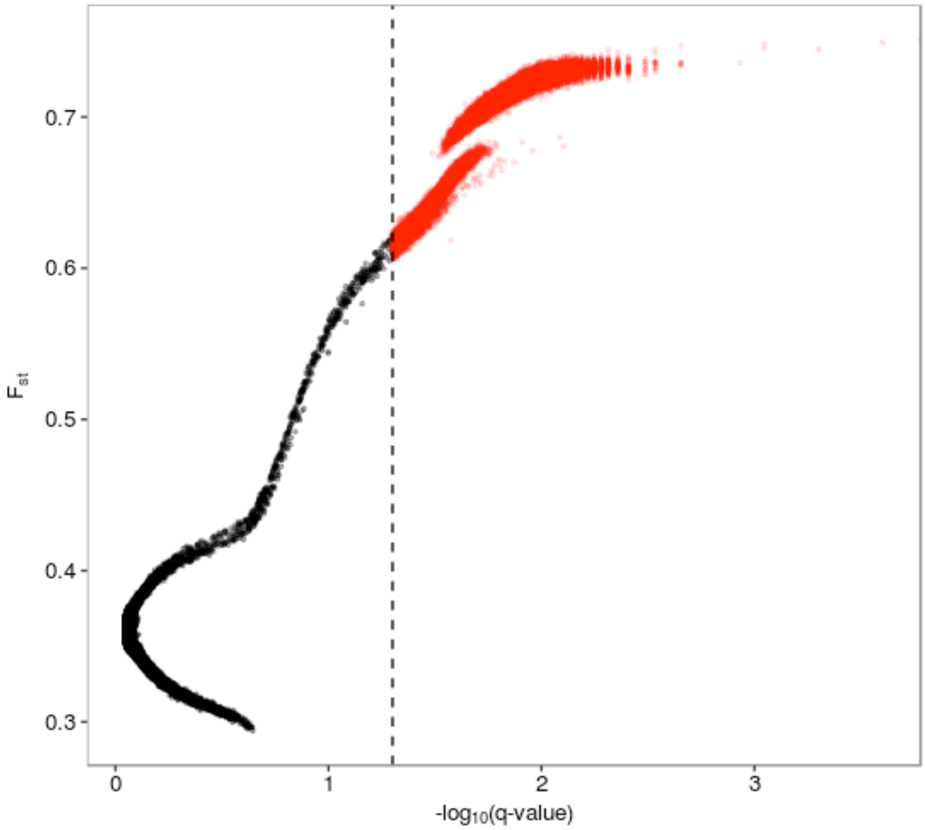


**Supplemental Figure 8 |** The overall $F_{st}$ estimates from the multi-way BayeScan analysis for SNPs plotted against their q-values. The SNPs below the q-value threshold of 0.05 (dashed vertical line) have been randomly sub-sampled to 10,000 representative SNPs for ease of plotting.

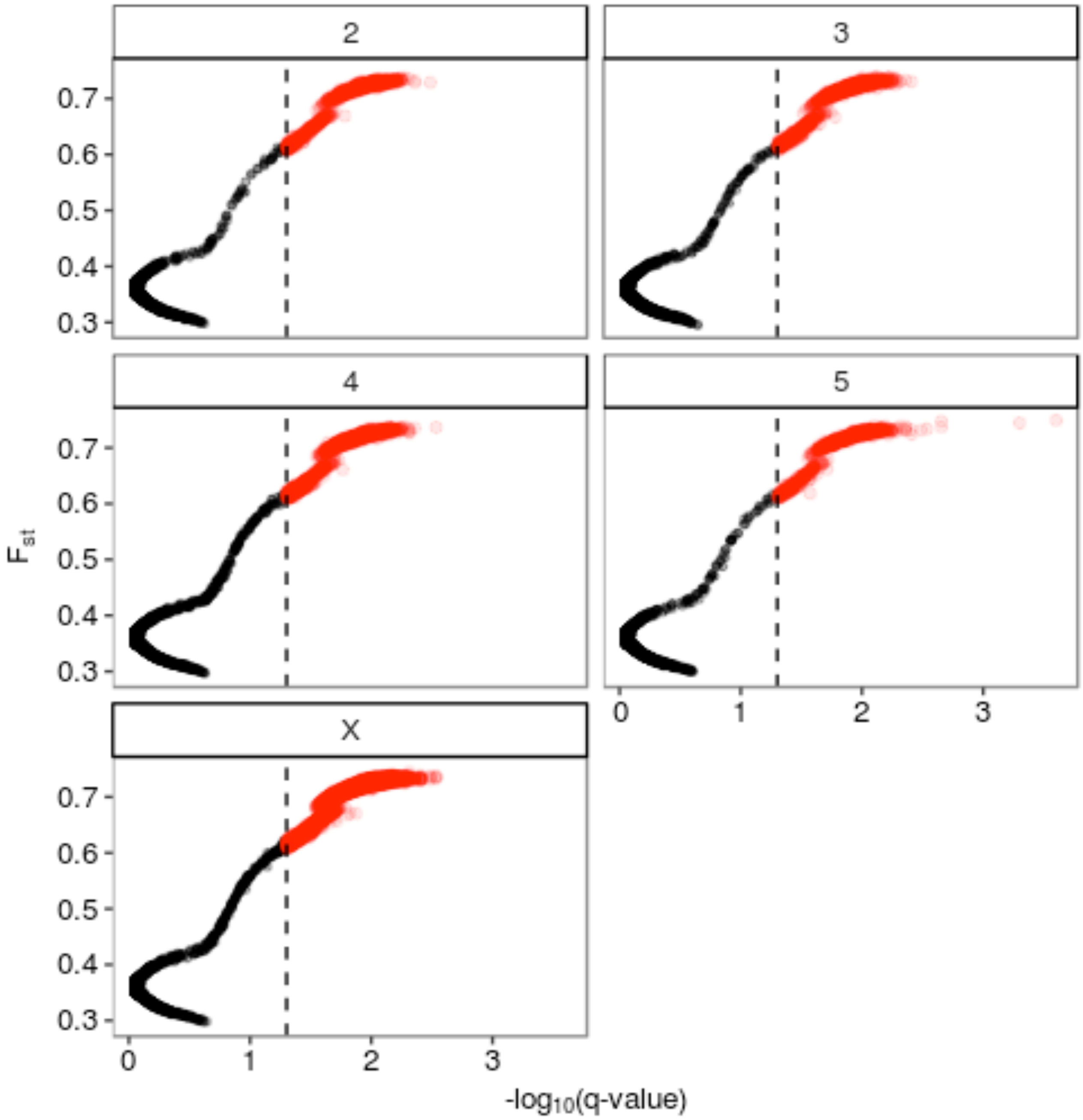


**Supplemental Figure 9 |** The $F_{st}$ estimates from the multi-way BayeScan analysis for SNPs on the mapped chromosomes plotted against their q-values. The SNPs below the q-value threshold of 0.05 (dashed vertical line) have been randomly sub-sampled to 10,000 representative SNPs for ease of plotting.

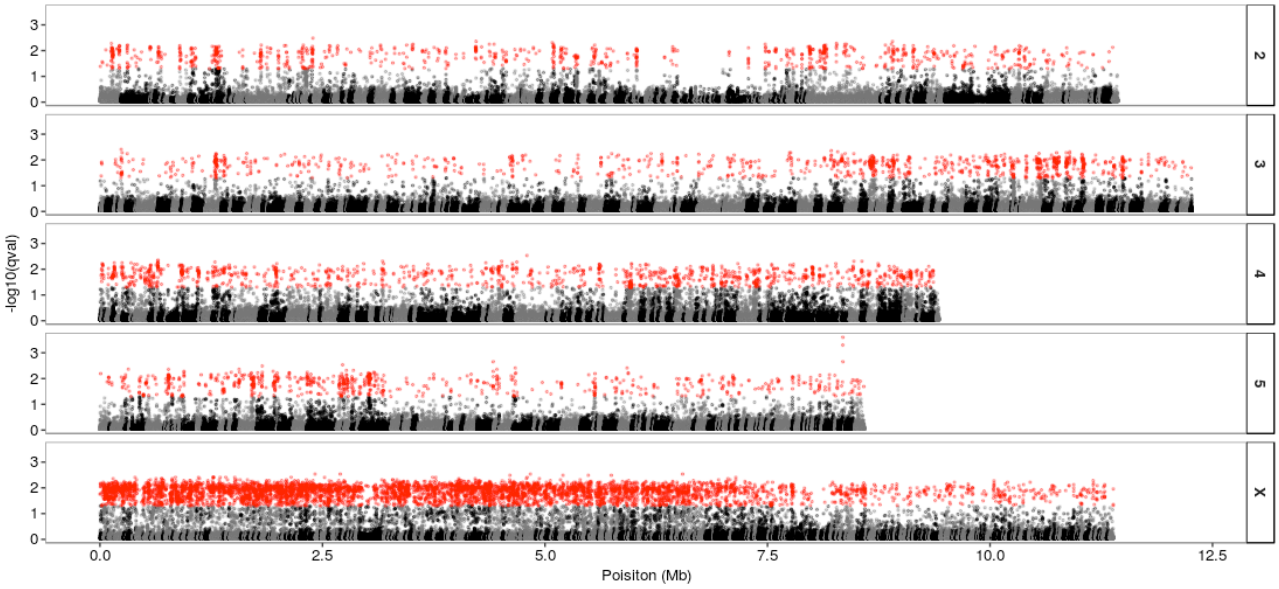


**Supplemental Figure 10 |** Manhattan plot of q-values from the multi-way BayeScan analysis for the SNPs on the mapped chromosomes. Red points denote SNPs which pass the 0.05 q-value FDR threshold. Alternating grey and black points denote different scaffolds which have been anchored to the chromosomes. The order of the mapped scaffolds is established but not their orientation.

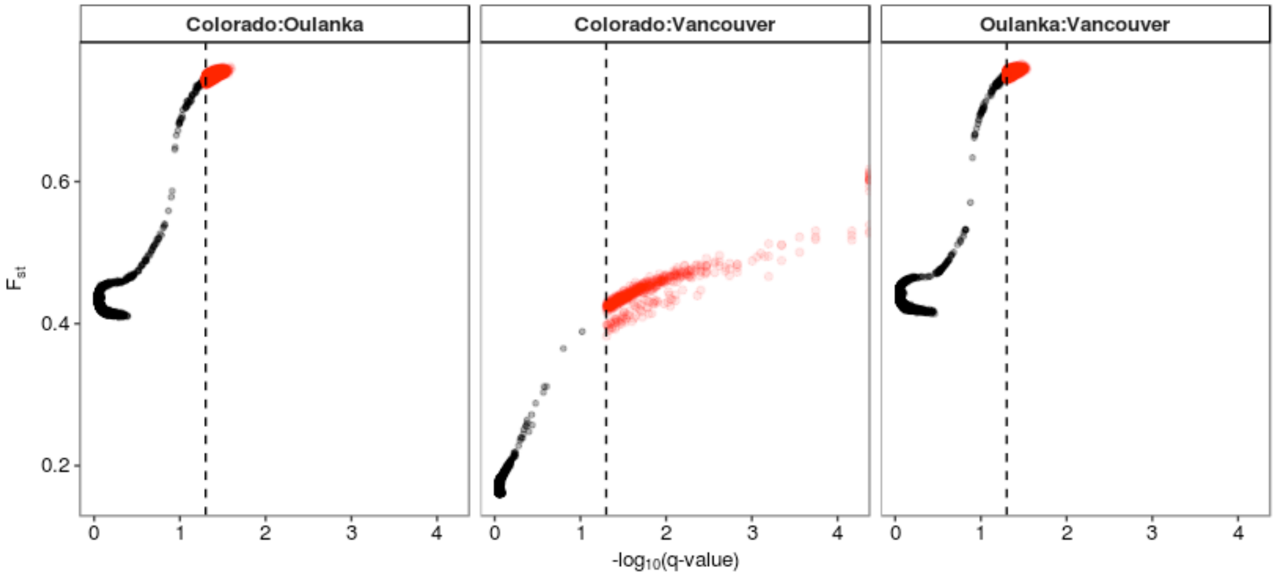


**Supplemental Figure 11 |** The overall $F_{st}$ estimates from the three pairwise BayeScan analyses for SNPs plotted against their q-values. The SNPs below the q-value threshhold of 0.05 (dashed vertical line in each panel) have been randomly sub-sampled to 10,000 representative SNPs for ease of plotting.

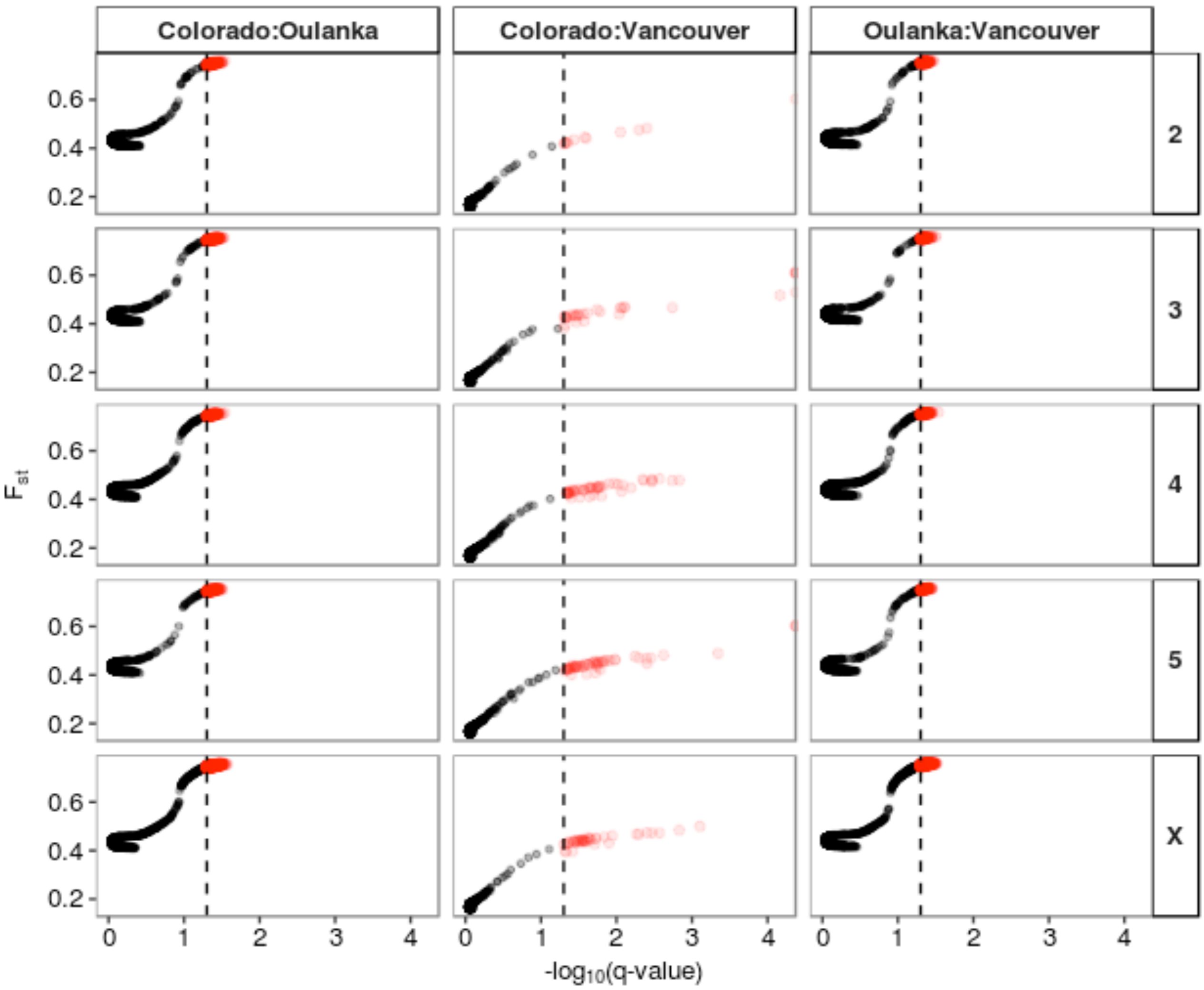


**Supplemental Figure 12 |** The $F_{st}$ estimates from the pairwise BayeScan analyses for SNPs on the mapped chromosomes plotted against their q-values. The panels in the vertical axis are the four main autosomes and the X-chromosome. The SNPs below the q-value threshold of 0.05 (dashed vertical line in each panel) have been randomly sub-sampled to 10,000 representative SNPs for ease of plotting.

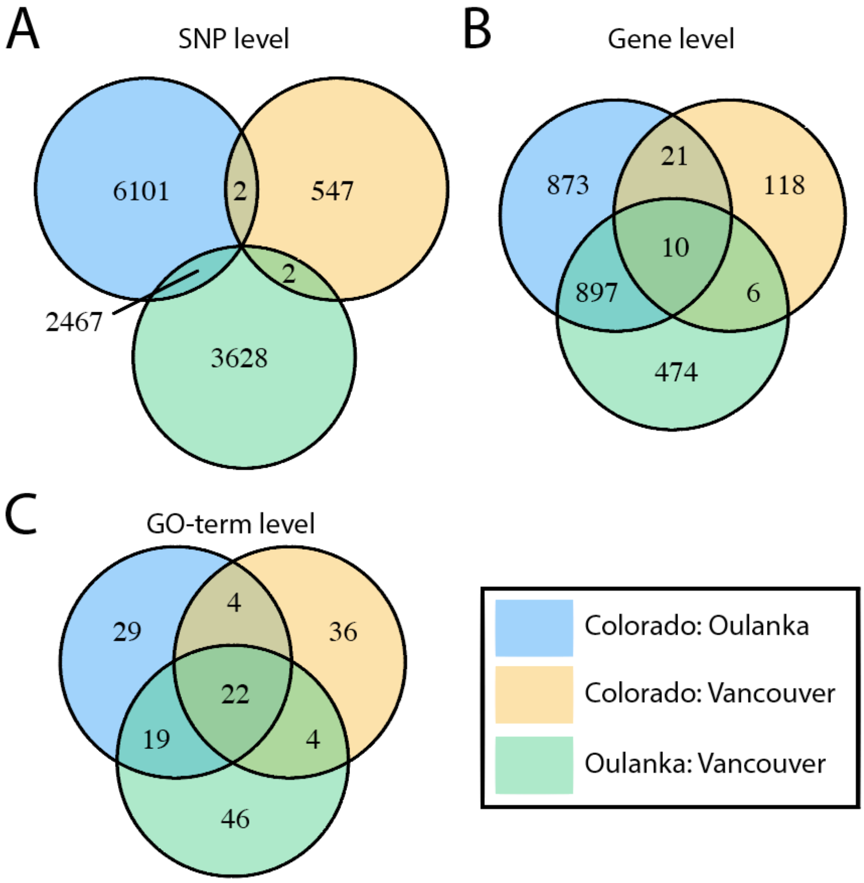


**Supplemental Figure 13 |** A) Overlap of significantly differentiated SNPs between populations B) Overlap of significantly differentiated genes between populations C) Overlap of enriched GO terms of significantly differentiated genes between populations
